# Metabolic switch in the aging astrocyte supported via integrative approach comprising network and transcriptome analyses

**DOI:** 10.1101/2022.01.31.478210

**Authors:** Alejandro Acevedo, Felipe Torres, Miguel Kiwi, Felipe Baeza-Lehnert, L. Felipe Barros, Dasfne Lee-Liu, Christian González-Billault

## Abstract

Dysregulated central-energy metabolism is a hallmark of brain aging. Supplying enough energy for neurotransmission relies on the neuron-astrocyte metabolic network. To identify genes contributing to age-associated brain functional decline, we formulated an approach to analyze the metabolic network by integrating flux, network structure and transcriptomic databases of neurotransmission and aging. Our findings support that during brain aging: 1) The astrocyte undergoes a metabolic switch from aerobic glycolysis to oxidative phosphorylation, decreasing lactate supply to the neuron, while the neuron suffers intrinsic energetic deficit by downregulation of Krebs cycle genes, including *mdh1* and *mdh2* (Malate-Aspartate Shuttle); 2) Branched-chain amino acid degradation genes were downregulated, identifying *dld* as a central regulator; 3) Ketone body synthesis increases in the neuron, while the astrocyte increases their utilization, in line with neuronal energy deficit in favor of astrocytes. We identified candidates for preclinical studies targeting energy metabolism to prevent age-associated cognitive decline.

## INTRODUCTION

Energy metabolism, essential for brain function, is one of the main processes dysregulated during brain aging (reviewed in Mattson and Arumugam, 2018; Pellerin and Magistretti, 1994). Although the brain constitutes only 2% of total body mass, it represents 20-25% of total body energy expenditure (Attwell and Laughlin, 2001; Bonvento and Bolaños, 2021), where most of it is used for re-establishing cation gradients after neurotransmission (Harris et al. 2012), a process mediated by sodium/potassium-ATPase pumps (Baeza-Lehnert *et al.* 2018; Erecinska & Silver 1994). To meet this high energy demand, the neuron and astrocyte form a two-cell metabolic network *(the neuron-astrocyte metabolic network)* with extensive metabolite exchange (Bonvento & Bolaños 2021; Bélanger et al. 2011). One example of metabolic exchange is the astrocyte-neuron lactate shuttle (ANLS) (Pellerin & Magistretti 1994). The astrocyte performs aerobic glycolysis, converts pyruvate into lactate, and then transports it to the neuron to fuel ATP synthesis via oxidative phosphorylation (Mächler et al. 2016; Zuend et al. 2020). The neuron-astrocyte metabolic network also performs the glutamate-glutamine cycle (GGC). In the GGC, astrocytes take up glutamate - the main excitatory neurotransmitter in the central nervous system-after neurotransmission. Inside the astrocyte, glutamate is converted into glutamine, shuttled back to the neuron, and re-converted into glutamate for a new neurotransmission cycle (Magistretti & Allaman 2015; Hasel *et al.* 2017; Schousboe 2019; Hyder et al. 2006). The ANLS, GGC, and the exchange of sodium and potassium constitute essential metabolic interactions between neurons and astrocytes, and they are closely related to energy metabolism. Indeed, energy availability is vital to ensure proper neurotransmission. However, during human brain aging, metabolism becomes dysregulated in the brain. Healthy aged human individuals display slower mitochondrial metabolism and glutamate-glutamine cycle neuronal flux (−28%) when compared with healthy young individuals. In comparison, astroglial mitochondrial flux is 30% faster (Boumezbeur *et al.* 2010). In rats, adult primary astrocyte cultures also display a higher mitochondrial oxidative metabolism when compared with astrocytes derived from young rats (Jiang & Cadenas 2014). To date, the only intervention demonstrated to extend lifespan in several model organisms is caloric restriction, a metabolic intervention where animal models are fed a diet consisting of 60-70% of the calorie intake in a regular diet (Mattison *et al.* 2017). This further supports the role of energy metabolism during aging. Metabolic challenges like the ketogenic diet (Newman *et al.* 2017; Roberts *et al.* 2017) and intermittent fasting that aim to mimic the metabolic state entered during caloric restriction have also been shown to extend lifespan and health-span (Dias *et al.* 2021). Furthermore, a phase II clinical trial using a fasting-mimicking diet improved metabolic health (Wei et al. 2017).

The complexity of brain aging is determined by the diversity and number of metabolic pathways that contribute to energy balance. The molecular mechanisms underlying age-associated dysregulation of brain energy metabolism remain mostly unknown. Complex systems -particularly metabolic pathways-are studied by modeling them as networks, which allows to simulate and probe complex phenomena, such as aging, in a computationally tractable and interpretable fashion (O’Brien et al. 2015; Zweig 2016). Here, we present a novel network-wise mapping complex interactions into a graph representation approach to discover energy-related genes in the neuron-astrocyte metabolic network that may contribute to brain aging. We used a genome-scale model of the neuron-astrocyte metabolic network (Lewis *et al.* 2010) and analyzed it using complementary flux and network-based methods. Flux-based methods allowed us to identify reactions critical for maintaining optimal neurotransmission.

On the other hand, network-based methods (centrality) searched for reactions that may modulate neurotransmission via network-wide effects. This analysis provided us with a set of genes (metabolic hub genes) that are key for neurotransmission in terms of flux distribution and network structure. Next, we determined which metabolic hub genes showed differential abundance associated with neurotransmission and/or brain aging in the neuron and/or astrocyte, thus getting a final set of genes called *differential hub genes* (DHG). These gene set represents a validation of network analysis contrasting numerical predictions with experimental data, including expected and novel results.

Functional annotation analysis of DHGs led to the following main findings: 1) Gene expression changes in both the neuron and astrocyte suggest an energetic deficit in the neuron, mainly by substantial downregulation of tricarboxylic acid (TCA) cycle in the aging neuron; 2) In line with the neuronal energy deficit, our results suggest that the aging astrocyte undertakes a metabolic switch from aerobic glycolysis to oxidative metabolism, where glucose is directed to CO_2_ instead of lactate; 3) Impaired branched-chain amino acid degradation in both the neuron and astrocyte, mainly supported by downregulation of the *dld* gene during aging. This gene encodes for a subunit of the branched-chain amino acid (BCAA) dehydrogenase complex, which catalyzes an early step in BCAA degradation; 4) Altered ketone body metabolism, where gene expression changes in the neuron agree with an increased synthesis during brain aging, while in the aging astrocyte *bdh1* is upregulated. This gene catalyzes the interconversion between acetoacetate and β-hydroxybutyrate, the main two ketone bodies required for ketone body utilization. These findings further support that energy metabolism is favored in the astrocyte, in detriment of neuronal energy supply; 5) Downregulation of genes associated with synaptic transmission in the neuron, including downregulation of Na/K-ATPase pumps in the aging neuron, and lower glutamate synthesis in both neuron and astrocyte; 6) Our results suggest that the aging neuron downregulates genes that supply the one carbon and tetrahydrofolate (THF) pool, which is required for the synthesis of the methylation precursor S-adenosylmethionine (SAM) and antioxidant glutathione synthesis. Instead, the aging astrocyte displays expression changes that agree with an increase in the THF pool available for glutathione synthesis as an antioxidant strategy, which is also in line with the metabolic switch into an oxidative metabolism in the astrocyte (which requires more antioxidation).

The genes identified here are valuable candidates for future studies to understand the molecular mechanisms of healthy brain aging and prevent brain age-associated failure using energy metabolism as a target. We also highlight how our approach significantly provides a robust and tractable number of final gene candidates for future studies, using an integrative analysis of the two-cell neuron-astrocyte metabolic network, which may be applied to other metabolic models.

## RESULTS

### Workflow Overview

To facilitate reading of the following sections, we provide an overview of the analyses performed in simple terms. We started by using a previously available genome-scale neuron-astrocyte (N-A) metabolic network model (Lewis et al., 2010), which included all metabolic reactions and transport events (each representing a *node* in the network) that occur in each cell across subcellular compartments, as well as transport between cell types. Genome-scale metabolic models are constructed using genome-wide gene expression data to only include nodes that are present in neurons and/or astrocytes (Lewis *et al.*, 2010) (Figure 1a). We used this N-A metabolic network to perform a *Flux Balance Analysis (FBA)* (Figure 1b) and a *Centrality Analysis* (Figure 1c). Broadly, the FBA calculates the extent to which the flux through each node in the N-A metabolic network should be modified for optimal achievement of the *metabolic objective*, which we defined as glutamatergic neurotransmission workload (*i.e.*, energy burden derived from neurotransmission). This is defined as the *optimal metabolic response*. The FBA identifies two types of nodes: *flux nodes*, which are those that most contribute to optimally achieving the metabolic objective of neurotransmission workload, and *sensitive nodes*, which are the key nodes exerting control over neurotransmission workload. Merging these two types of nodes yielded the list of *optimal nodes*, where each node has been previously associated with specific genes. We defined *optimal genes* as the list of genes associated with optimal nodes.

**Figure 1.**
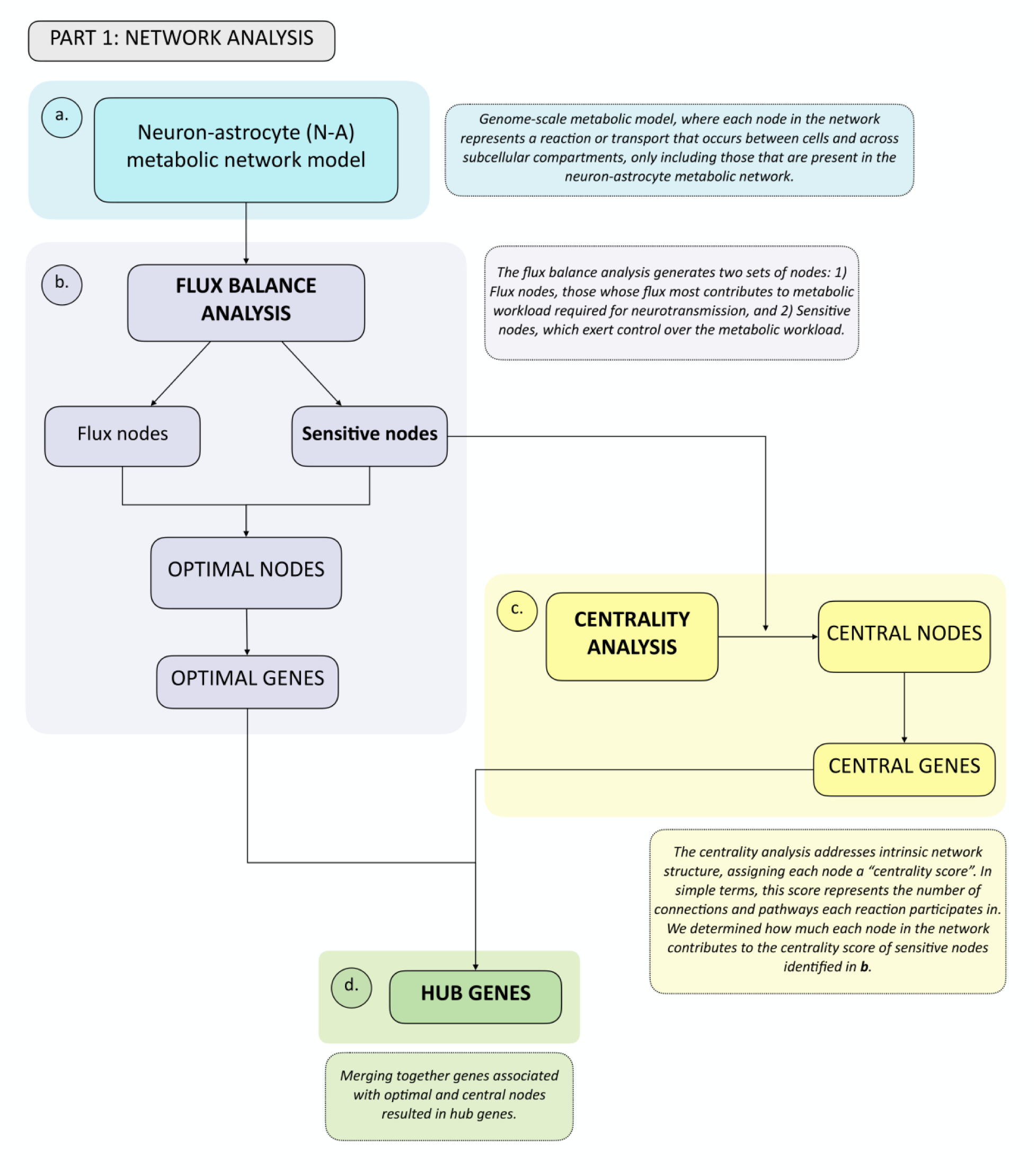
Summary flowchart of network analyses depicting how optimal and central genes were identified, which merged together form the hub genes group. **a.** A genome-scale metabolic model from Lewis et al., 2010 was used. This network was analyzed first using **b.** Flux Balance Analysis, from which Flux and Sensitive Nodes were identified. Merging these two node lists yielded Optimal Nodes, from which Optimal Genes were identified. Sensitive Nodes were then analyzed using **c.** Centrality Analysis, which allowed identifying Central Nodes, from which Central Genes were identified. Merging the list of Optimal and Central Genes produced the Hub Genes list **(d)**. See boxes in dashed lines for the explanation of each type of analysis.

The FBA was followed by a centrality analysis, which analyzes intrinsic network structure, and is therefore independent of flux. In a centrality analysis, each node in the network has a *centrality score*, which, largely, represents the number of connections and pathways each node participates in. We calculated how the removal of each individual node in the network affected the centrality score of sensitive nodes identified in the previous step, as those represent the ones that exert control over the metabolic objective. Nodes significantly altering the centrality of sensitive nodes were defined as *central nodes*, from which the list of *central genes* was obtained. By merging optimal and central node lists we obtained the list of *hub genes* (Figure 1d), which represent the genes that most affect glutamatergic neurotransmission workload, and therefore play a key role in N-A metabolic network function.

Having identified hub genes that play key roles in the N-A metabolic network, we next determined which of these were differentially expressed, *i.e.*, up- or downregulated after neurotransmission and/or brain aging in the neuron and/or astrocyte (2a). To achieve this, we used previously available transcriptomic databases for neurotransmission (2b) and brain aging (2c) (Consortium, 2020; Hasel et al., 2017). The last step in gene selection identified hub genes that were differentially expressed during neurotransmission and/or brain aging (see shaded area in Venn diagram, 2d) defined as differential hub genes (DHG) (2e). This curated group of genes represents those that most contribute to achieving glutamatergic neurotransmission workload. The ultimate goal of this integrative analysis was to identify genes and pathways important for neurotransmission, which fail during brain aging, thus constituting candidates to explain age-associated cognitive decline. To achieve this, the final step was a KEGG *pathway enrichment analysis* of DHG, which allowed us to identify the predominant metabolic pathways.

### Flux-based analysis identifies optimal nodes in the neuron-astrocyte network required for glutamatergic neurotransmission workload

Regarding the FBA (Figure 1b), in this analysis we defined three sub-objectives that represent key processes required for achieving neurotransmission workload: 1) The astrocyte-neuron lactate shuttle (ANLS), 2) The glutamate-glutamine cycle (GGC), and 3) Sodium removal by Na/K-ATPase pumps (Figure 3a-c). The FBA therefore determined how to optimize flux through these three processes by identifying flux and sensitive nodes (Figure 1b). Furthermore, for the results to be biologically coherent, we used experimentally determined flux values during neurotransmission as *constraints*. These were the neuronal and astrocytic glucose and oxygen consumption rates, and neuronal ATP maintenance rate, reported by Fernandez-Moncada et al. 2018 and Baeza-Lehnert et al., 2018. Also, metabolite *steady state* was imposed as a constraint. This means that intracellular metabolite concentration levels remain constant under neurotransmission (see Supplemental Theoretical Framework Section 1.2 on how this is relevant for the analysis).

Figures 2d-g depict fluxes previously associated with the metabolic sub-objectives ANLS, GGC and Na/K-ATPase pumps in *phenotypic phase planes* (PhPPs), where non-zero slopes can be observed (see Methods section Phenotypic Phase Plane Analysis for details). These allowed validating that each sub-objective is dependent on oxygen and glucose uptake rates, which is a hallmark of brain metabolism. The optimal flux that maximizes each metabolic sub-objective is shown as a red-filled circle in each PhPP (Figures 2d-g). Figure 2d shows that the calculated optimal neuronal sodium efflux associated with removal through Na/K-ATPase pumps was 350 uM/s. Also, Figure 2e shows that lactate efflux from the astrocyte was 6.913 uM/s, and Figure 2f that vesicle-mediated export of glutamate from the neuron was 4.138 uM/s (influx into the complementary cell and other relevant fluxes are shown in Supplemental Table S1). Furthermore, from Figure 3g it is possible to assume that the optimal solution is unique since it is located on a vertex. In addition, the optimal metabolic response was associated with complete (aerobic) glucose oxidation. In this sense, six oxygen molecules oxidized one glucose molecule (Supplemental Figure S1a), while ATP yield was close to 27.5 ATP molecules per glucose molecule (Supplemental Figure S1b). Of note, it is possible that this last yield was lower than the theoretical one due to flux to other pathways such as the pentose phosphate pathway and reactions that exit the tricarboxylic acid cycle (TCA), *e.g.*, glutamate synthesis and the malate-aspartate shuttle (MAS). Furthermore, in line with what Baeza-Lehnert et al. (2018) reported, we observed flux coupling between ATP demand from the sodium ATPase pump and ATP supply from oxidative phosphorylation in neurons (Figure S2). Overall, our model was mathematically consistent and agreed with the biology of neurons and astrocytes undergoing neurotransmission.

**Figure 2.**
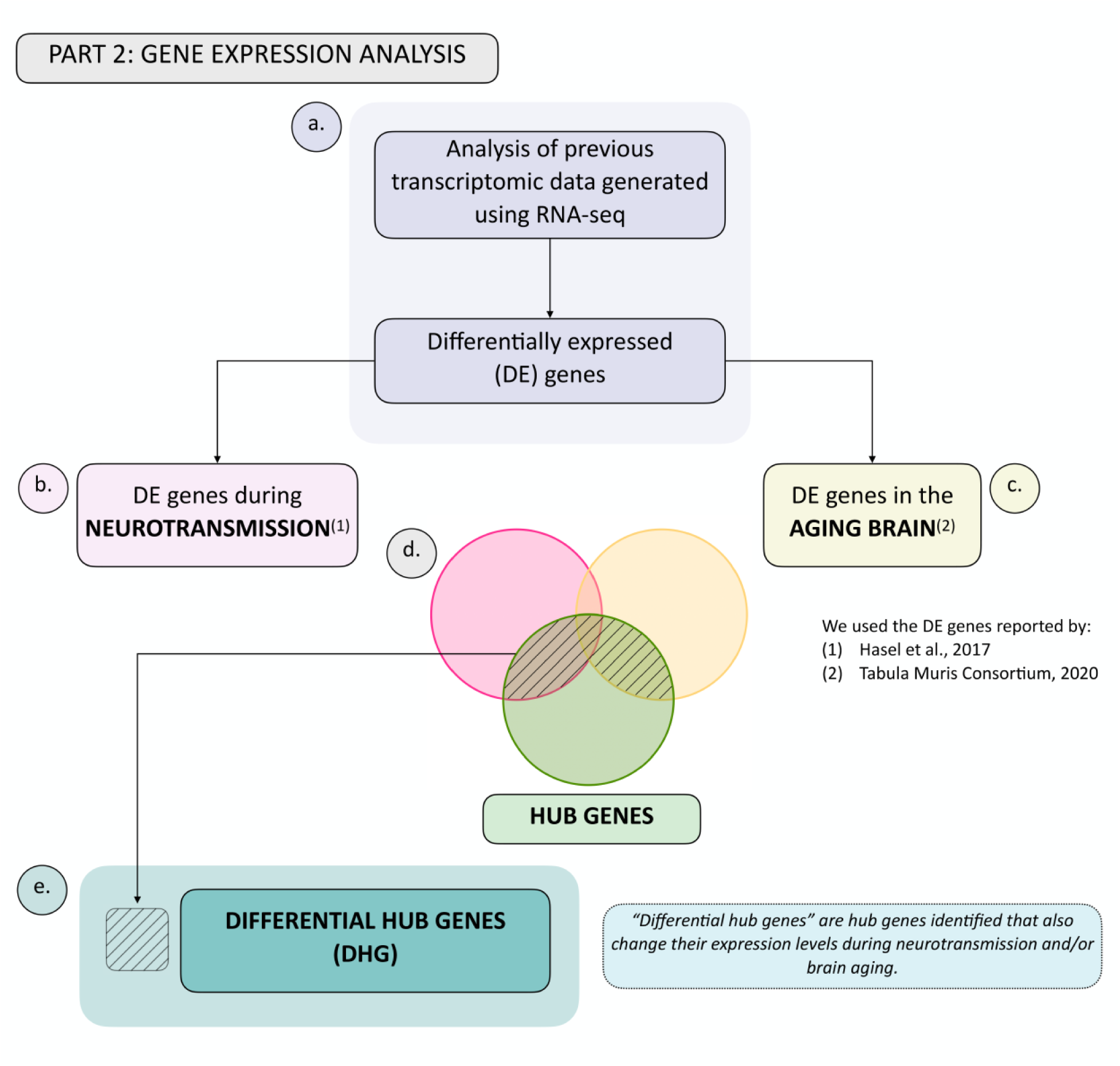
Summary flowchart of integration of hub genes with transcriptomic data generated during neurotransmission and brain aging. **a.** Transcriptomic data during neurotransmission (Hasel et al., 2017) and aging (Tabula Muris Consortium, 2020), reporting differentially expressed genes during each process in the neuron and/or astrocyte was obtained. This allowed us to obtain a list of differentially expressed (DE) genes in both cell types during **b.** neurotransmission and/or **c.** brain aging. **d.** Venn diagram showing common genes: 1) Between DE genes during neurotransmission and hub genes (pink and green sets); 2) Between DE genes during brain aging and hub genes (yellow and green sets), and 3) The intersection between all three gene groups (pink, yellow and green sets). **e.** The differential hub genes (DHG) list is shown in d. in the shaded area.

**Figure 3.**
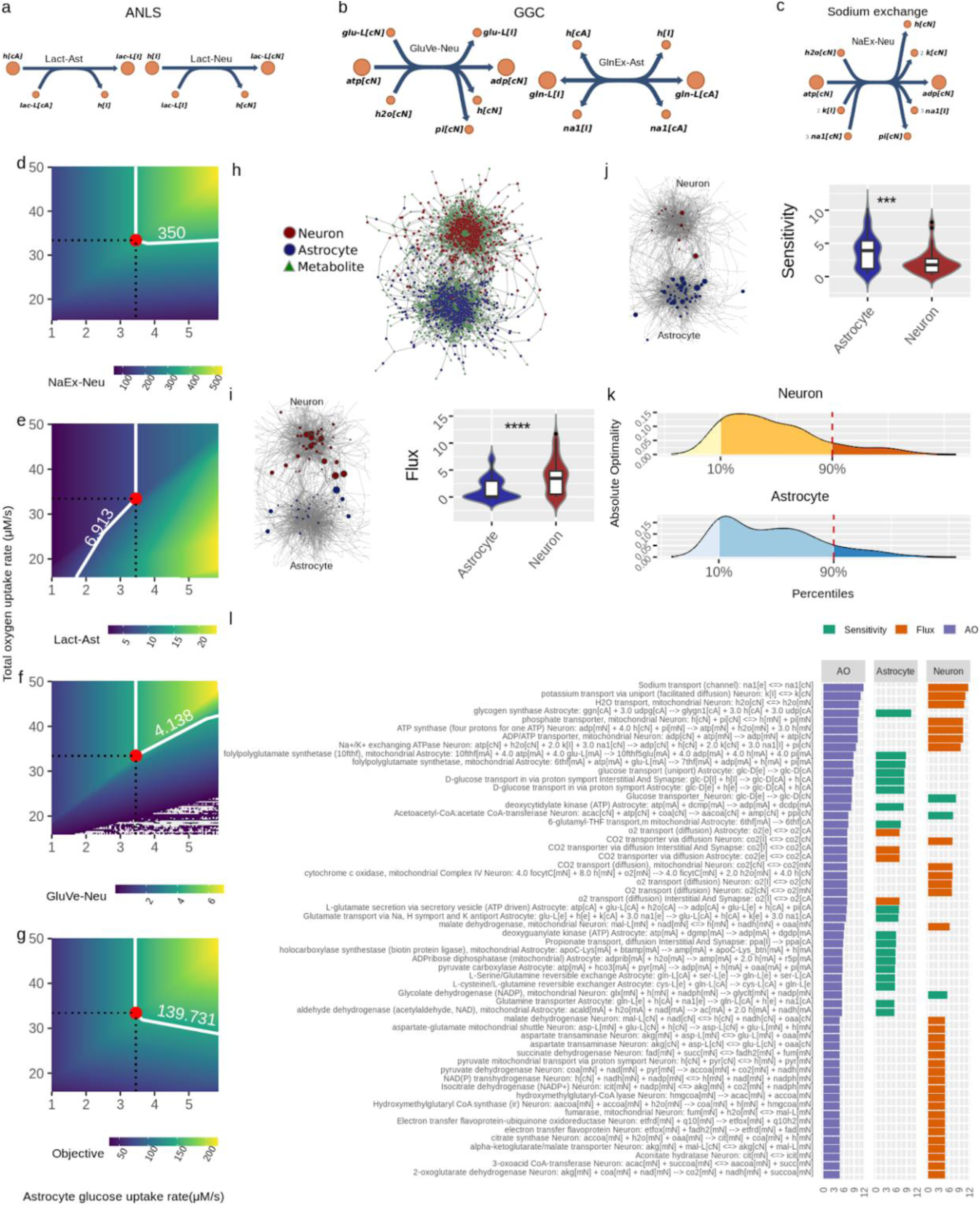
Identification of optimal nodes using Flux Balance Analysis in the neuron-astrocyte metabolic network suggests division of labor between the neuron and astrocyte in response to neurotransmission workload. **a-c.** Reactions considered in the metabolic objective; here, metabolite names correspond to the same as in the model reported by Lewis et al. (2010). a. Fluxes associated with the Astrocyte-Neuron Lactate Shuttle (ANLS); *left side*: Lactate efflux from astrocyte to the interstitial space (Lact-Ast); *right side*: Lactate from the interstitial space entering neurons (Lact-Neu). **b.** Fluxes related to the Glutamate-Glutamine Cycle (GGC); *left side*: vesicle-exported glutamate from neuron (GluVe-Neu); *right side*: glutamine excretion from astrocyte (GlnEx-As). c: Neuronal sodium efflux associated with its removal via sodium ATPase pump. **d-g.** Phenotypìc phase planes are shown as two-dimensional color maps. Here, the Flux Balance Analysis (FBA) solution is represented by the red-filled circle, while all fluxes shown correspond to micromolar per second (uM/s). A white piece-wise line depicts the specific contour level of the solution. **h:** The neuron-astrocyte metabolic network is represented as a bipartite network; here, node shape (circle or square) denotes the partition where it belongs, i.e., reaction or metabolite. **i.** *left side,* flux values distribution in each cell; *right side*: the bipartite network presented in **(h)** showing node size proportional to absolute flux. **j.** *left side,* sensitivity values distribution in each cell; *right side*: the bipartite network presented in **(h)** showing node size proportional to absolute sensitivity. **k.** Distribution of the Absolute Optimality values in neuron and astrocyte, the 90 percentile is highlighted by a red dashed line. This line depicts the cutoff over which a reaction was classified as an optimal metabolic reaction. **l.** Optimal metabolic reactions (descending order) sorted by their Absolute Optimality and presented alongside their flux and sensitivity.

In addition to fluxes, the optimal metabolic response is shaped by *sensitivity*, which is equally relevant to flux in the FBA (Acevedo et al., 2014; Acevedo et al., 2017). Sensitivity values inform the extent to which a change in any given reaction modifies the optimal metabolic response. We calculated sensitivities and, together with fluxes, determined how they distributed throughout the neuron-astrocyte metabolic network. Interestingly, high-flux reactions were mostly neuronal (Figure 2i), while high-sensitivity reactions were mainly astrocytic (Figure 2j). This cellular separation among flux and sensitivity suggests neurotransmission sets up fluxes in neurons, and sensitivities in astrocytes. Next, we combined the flux and sensitivity of each node into a single quantity called *Absolute Optimality* (AO) (see Methods section Absolute Optimality for details). The AO informed us about the involvement any given node has in the achievement of the optimal response. All nodes that had an AO above the significant threshold were considered *optimal nodes* (Figures 1b and 2k). Figure 2l shows fluxes and sensitivities of optimal nodes separated by cell type and sorted in descending order for AO.

Taken together, the optimality analysis suggests a division of labor between neurons and astrocytes in response to neurotransmission workload. Here, the execution, represented by flux, is allocated to neurons, while control, represented by sensitivity, is executed by astrocytes.

### Analysis of network structure based on sensitive nodes further supports the division of labor between the neuron and astrocyte in the network

We further analyzed the N-A metabolic network to enrich our analysis, by performing a centrality analysis (Figure 1c). While part of aging-derived damage to brain metabolism may reside in fast stationary events such as those represented by FBA results, much of aging deterioration may occur in non-steady state long-term events. Intrinsic network structure allows identifying long-term phenomena beyond steady state and short timescales (see Methods *Modeling rationale*). As mentioned before, the centrality score of a node represents how connected the node is in the network. We calculated the extent to which each node in the network, when removed, affected the centrality of sensitive nodes identified in the previous step (see Methods *Absolute Centrality Contribution*). Four complementary centrality metrics were employed to ensure analysis robustness; thus, each reaction was associated with four quantities. These accounted for how much a given reaction contributes to the centrality of the sensitive nodes and were denominated *centrality contributions*. As can be observed in Figure 4a, in astrocytes centrality contributions tended to be positive, while in neurons it was mostly negative. This result indicates that astrocytic nodes tend to increase the centrality of the sensitivity set, while neuronal nodes tend to decrease it. This finding suggests opposite and complementary roles between cells.

**Figure 4.**
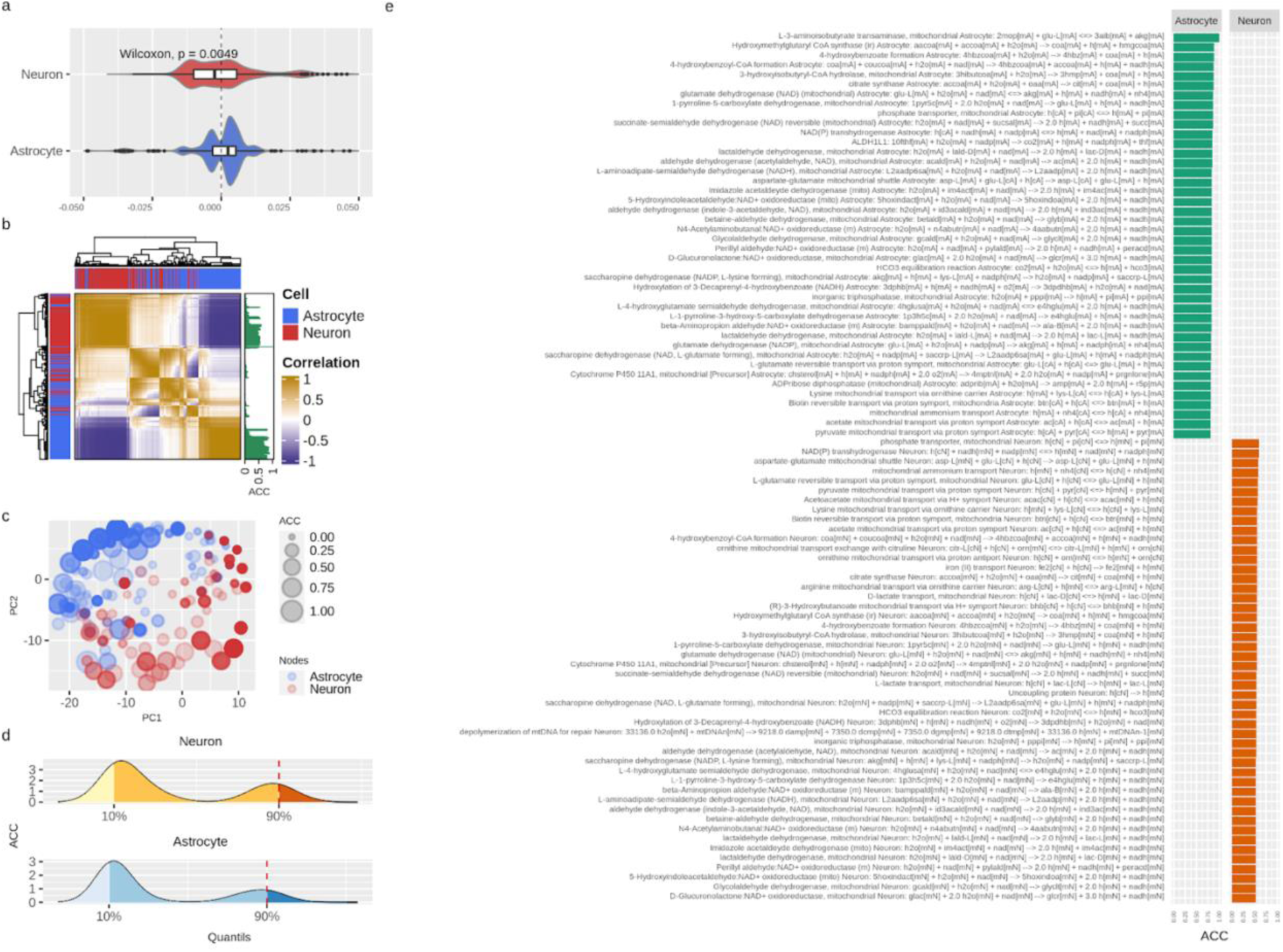
Centrality-based analysis of the neuron-astrocyte metabolic network further supports the division of labor between the neuron and astrocyte. **a.** Distributions, separated by cell, of the contributions of each reaction to the centrality of the sensitivity set. **b.** Unsupervised hierarchical clustering of the pairwise correlations between the contributions of each reaction to the centrality of the sensitivity set. The Absolute Centrality Contribution per reaction (ACC) is shown on the right-hand side of the heatmap. **c.** Dimensionality reduction via Principal Component Analysis (PCA) of the pairwise correlations between the contributions of each reaction. **d.** Distribution of ACC in the neuron (top) and astrocyte (bottom), here, the red dashed line by the 90% percentile indicates the cutoff over which reactions were considered central metabolic reactions. **e.** ACC values for the central metabolic reaction separated by cell.

In Figure 4b, this behavior was confirmed via unsupervised clustering of the correlations between the centrality contributions of each node (see Methods section for details on this procedure). Here we see that centrality contributions from the same cell are clustered together. The latter was also confirmed via dimensionality reduction, where the 2-dimensional distribution of the centrality contributions also resembled the two-cell structure (Figure 4c). Next, we aggregated the four centrality contributions into a single index which was a normalized and absolute value representing the capacity of a node to change the centrality of sensitivity nodes. We called this index *Absolute Centrality Contribution* (ACC). The ACC for each reaction is shown on the right-hand side of the heatmap in Figure 4b (see the column with green bars). Finally, the nodes with the last tenth percentile of the ACC values from each cell were categorized as *central nodes* (Figure 4d). Interestingly, the astrocyte concentrated the highest ACC values (Figure 4e). Merging optimal and central genes resulted in the hub genes list, which represent the genes with the highest probability to affect or control the N-A metabolic network in achieving glutamatergic neurotransmission workload.

As a whole, positive centrality contributions in the astrocyte and negative in the neuron, along with the predominantly high ACC of the astrocyte suggest well-differentiated roles for the neuron and astrocyte. These results are in the same line with those obtained by the FBA supporting the division of labor between the two cells.

### Identification of hub genes differentially regulated during neurotransmission and/or brain aging

Previously identified hub genes represent the scaffolding required for achieving glutamatergic neurotransmission, and among these, we sought to identify which were also differentially expressed during neurotransmission and/or brain aging. Disruption of these genes should lead to subpar neurotransmission workload, and therefore provide molecular insights into aging-associated brain functional decline. We denominated this group differential hub genes (DHG). To achieve this, we determined which of these were differentially expressed, *i.e.*, up- or downregulated after neurotransmission and/or brain aging in the neuron and/or astrocyte (Figure 2a). We used available transcriptome databases for neurotransmission (Figure 2b) and brain aging (Figure 2c) (Consortium, 2020; Hasel *et al.*, 2017) (see shaded area in Venn diagram, Figures 2d and 2e).

On the one hand, the neurotransmission database reported transcriptomic changes occurring in neurons and astrocytes grown in a mixed culture setting, before and after neurostimulation, followed by RNA-seq (Hasel *et al.*, 2017). The authors reported 4441 genes with differential abundance in the neuron and 1307 in the astrocyte (fold-change, FC ≥1.3 or ≤0.77 and padj-SSS-value < 0.05). On the other hand, the brain aging database was generated using single-cell RNA sequencing to obtain the age-coefficient for each gene, which is equivalent to the fold-change of each gene when comparing neurons and astrocytes from aged and young mouse brains (Consortium, 2020). This study reported 5415 differentially abundant genes in neurons and 1294 in astrocytes when comparing 1-3 months old with 18-30 months old mice (age-coefficient threshold at 0.005 reported by authors as equivalent to a 10%-fold change and an FDR threshold of 0.01).

The differentially expressed genes reported in these databases were then cross-referenced to the hub genes identified in the network analyses, resulting in DHG (Figures 2d and 2e). In response to neurotransmission, we found 53 DHG in the neuron and 14 DHG in the astrocyte. While for brain aging, we found 73 in the neuron and 26 in the astrocyte.

### Differential hub genes in the neuron suggest a metabolic deficit and impaired synaptic transmission during brain aging

We performed a pathway enrichment analysis using the KEGG pathway database, followed by manual curation to obtain a functional characterization of DHG in neurotransmission and brain aging (see Methods for manual curation criteria). Figure 5 shows KEGG pathways enriched in neuronal DHG during neurotransmission (Figures 5a to 5f) and brain aging (Figures 5a’ to 5f’), where node colors indicate up (red nodes) or downregulation (blue nodes) during each process.

**Figure 5.**
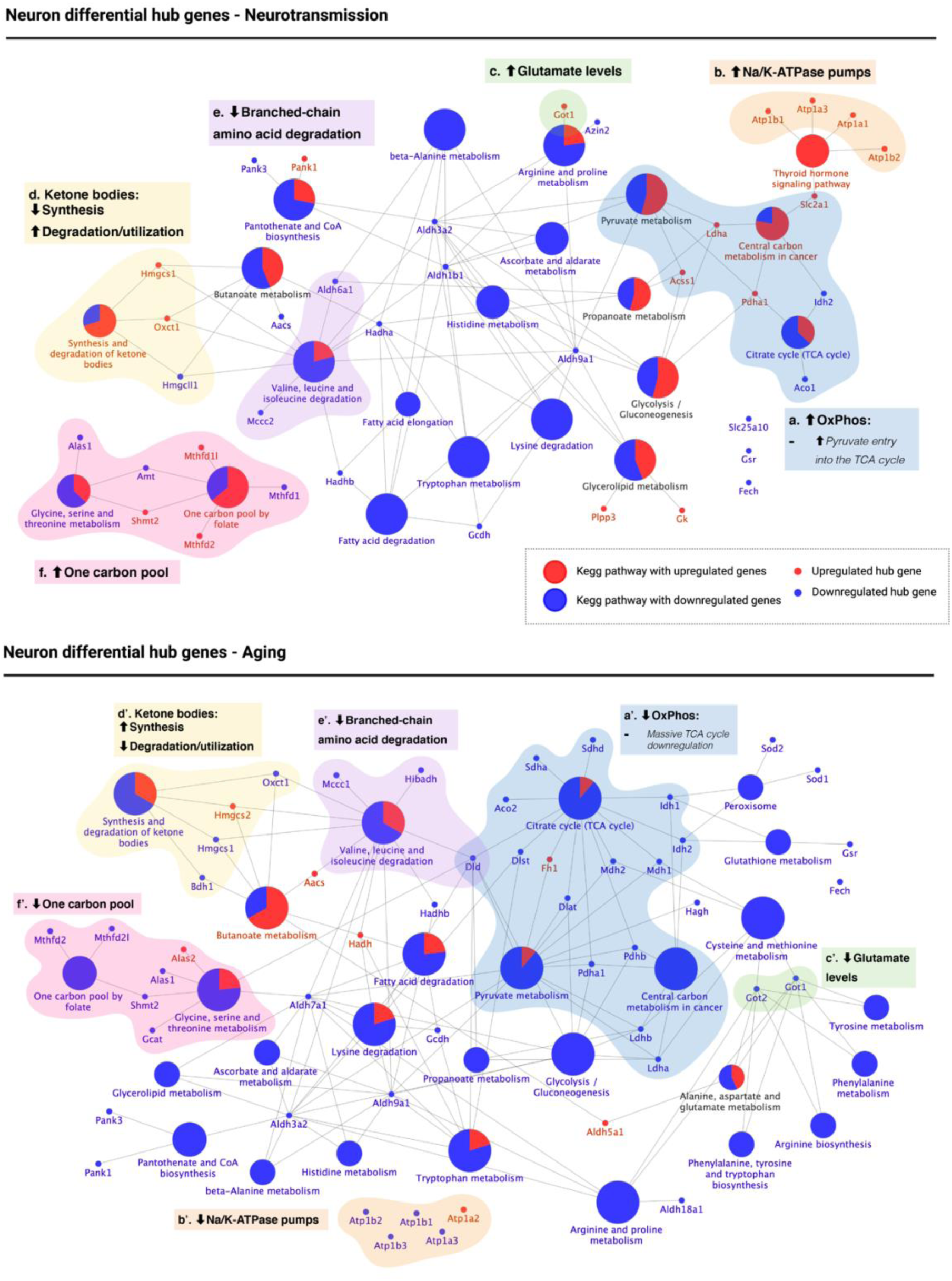
KEGG pathway enrichment of differential hub genes reveals that the aged neuron displays energetic deficit, dysfunctional neurotransmission, decreased branched-chain amino acid degradation and utilization of ketone bodies, and decreased one-carbon pool levels. KEGG pathway enrichment of differential hub genes was followed by manual curation of associated genes. The results are shown for neurotransmission (top panel) and aging (bottom panel). Oxidative phosphorylation (OxPhos, blue): high OxPhos levels during neurotransmission **(a)** but low OxPhos levels during aging **(a’)**. Synaptic transmission: upregulated Na/K-ATPase pumps (orange) and glutamate synthesis (green) suggest active re-establishment of cation gradients **(b)** and high glutamate levels **(c)**. The opposite was observed during aging **(b’-c’)**. 3) Ketone body metabolism (yellow): decreased synthesis and increased degradation/utilization during neurotransmission **(d)**, with the opposite observed during aging **(d’)**. 4) Branched-chain amino acid (BCAA) degradation (purple): while differential hub genes involved in the degradation of BCAA were found downregulated during both neurotransmission **(e)** and aging **(e’)**, *dld*, which encodes for a subunit of BCAA-decarboxylase, an early step in the degradation of all three BCAA was only downregulated during brain aging. 5) One carbon pool (pink): differential hub gene expression associated with one-carbon metabolism suggests high levels of one-carbon pool intermediates during neurotransmission **(f)** but low during aging **(f’)**.

We identified five main biological processes with different regulation when comparing neurotransmission and brain aging. The first group contained DHG associated with central energy metabolism associated with KEGG pathways “Pyruvate metabolism”, “Citrate cycle (TCA cycle)” and “Central carbon metabolism in cancer” (Figures 5a and 5a’, blue). This last pathway was included because metabolic changes observed in cancer, such as the Warburg effect, also occur in the brain (Barros et al., 2020). During neurotransmission, we observed upregulation of *acss1*, a mitochondrial enzyme that synthesizes acetyl-CoA from acetate, and of *pdha1*, which encodes for a subunit of the pyruvate dehydrogenase complex (PDC) (Figure 5a, blue). Upregulation of both enzymes agrees with increased acetyl-CoA levels and therefore suggests increased TCA flux, which would lead to high levels of oxidative phosphorylation. Instead, during aging, we observed downregulation of most genes involved in the three KEGG pathways mentioned above (except for *fh1*, which was upregulated). Notably, most of these DHG downregulated during neuronal aging participate in the TCA cycle. Plus, we found upregulation of three genes encoding for PDC subunits: *pdha1*, *pdhb,* and *dld*. These changes also suggest that acetyl-CoA entry into the neuronal TCA cycle and TCA cycle activity are impaired in the aged brain.

The second group was associated with synaptic activity, including a cluster of Na/K-ATPase pumps (Figures 3b and 3b’, orange) and enzymes that catalyze glutamate synthesis (Figures 5c and 5c’, green). During neurotransmission, they were upregulated, while in brain aging, they were downregulated except for *atp1a2*. Na/K-ATPase pumps are required to re-establish ion gradients after neurotransmission to allow the following cycle of synaptic activity. At the same time, glutamate is the primary excitatory neurotransmitter, for which these results agree with synaptic activity dysregulation during brain aging, with *got1/2* as DHG regulating glutamate levels.

The third group corresponds to the “Synthesis and degradation of ketone bodies” pathway (Figures 5d and 5d’, yellow). During neurotransmission, *hmgcs1*, encoding for the cytosolic form of 3-hydroxy-3-methylglutaryl-CoA synthase 1 was upregulated while *hmgcll1* (3-hydroxymethyl-3-methylglutaryl-CoA lyase like 1) was downregulated. *Hmgcs1* catalyzes the formation of HMG-CoA, which is further converted into mevalonate for cholesterol synthesis (as opposed to the mitochondrial isoform *hmgcs2*, which catalyzes the first irreversible step in ketogenesis using the same substrates as *hmgcs1*). Instead, *hmgcll1*, which catalyzes the second irreversible step in ketogenesis and is downregulated, and *oxct1*, which catalyzes the interconversion between acetoacetyl-CoA and acetoacetate and was upregulated. These results suggest the downregulation of ketone body synthesis during neurotransmission while favoring an increased degradation or utilization (Figure 3d). In contrast, during neuronal aging, it is *hmgcs2* which is upregulated (ketone body synthesis mitochondrial isoform), while *hmgcs1* is downregulated. In addition, *oxct1* and *bdh1* are also downregulated (Figure 5d’). *Bdh1* catalyzes the interconversion between acetoacetate and beta-hydroxybutyrate, the two main ketone bodies. Therefore, the downregulation of *oxct1* and *bdh1* suggest a decrease in ketone body turnover in the aged neuron.

The fourth group was associated with the “Valine, leucine, and isoleucine degradation” pathway, and therefore refers to branched-chain amino acid (BCAA) degradation (Figures 5e and 5e’, purple). While enzymes associated with BCAA degradation were downregulated during both neurotransmission and brain aging in the neuron, *dld*, which encodes for a subunit of the BCAA decarboxylase and thus catalyzes one of the first steps of the degradation of all three BCAA was only downregulated during brain aging (Figure 5e’), supporting downregulation of BCAA degradation during neuronal aging but possibly not during neurotransmission.

Finally, the fifth group was associated with regulating one-carbon pool levels (Figures 5f and 5f’, pink), including pathways “Glycine, serine and threonine metabolism” and “One carbon pool by folate.” During neurotransmission, glycine degradation enzymes *alas1* and *amt* were downregulated, while *shmt2*, which feeds the one-carbon pool by producing 5,10-methylenetetrahydrofolate was upregulated. Also, 2 out of 3 enzymes involved in the metabolism of one-carbon pool intermediates were upregulated (Figure 5f, pink). However, all enzymes (except for *alas2*) associated with these two pathways were downregulated during neuronal aging, suggesting a decrease in the one-carbon pool. We present a summary of all these changes in Table 1.

**Table 1.**
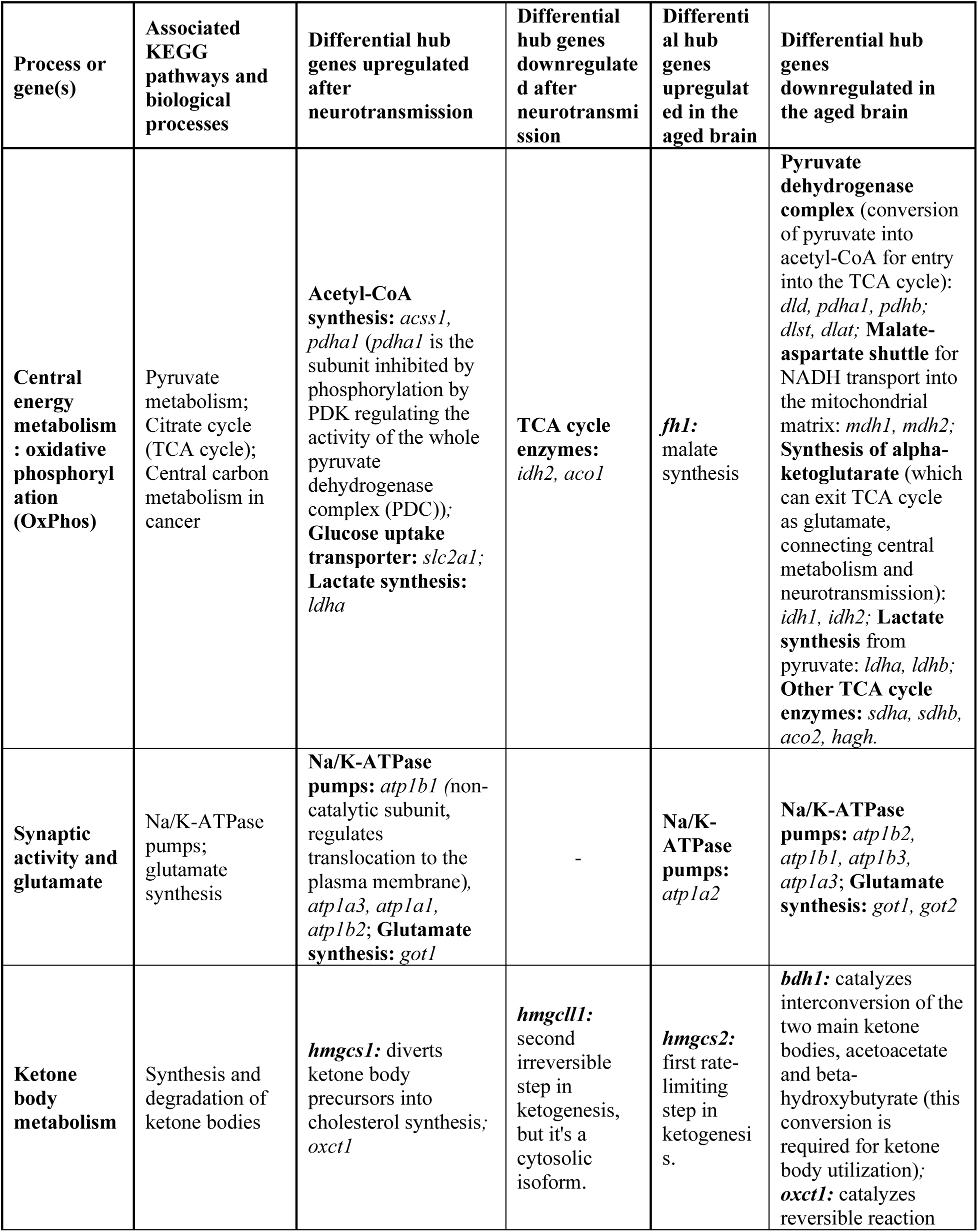
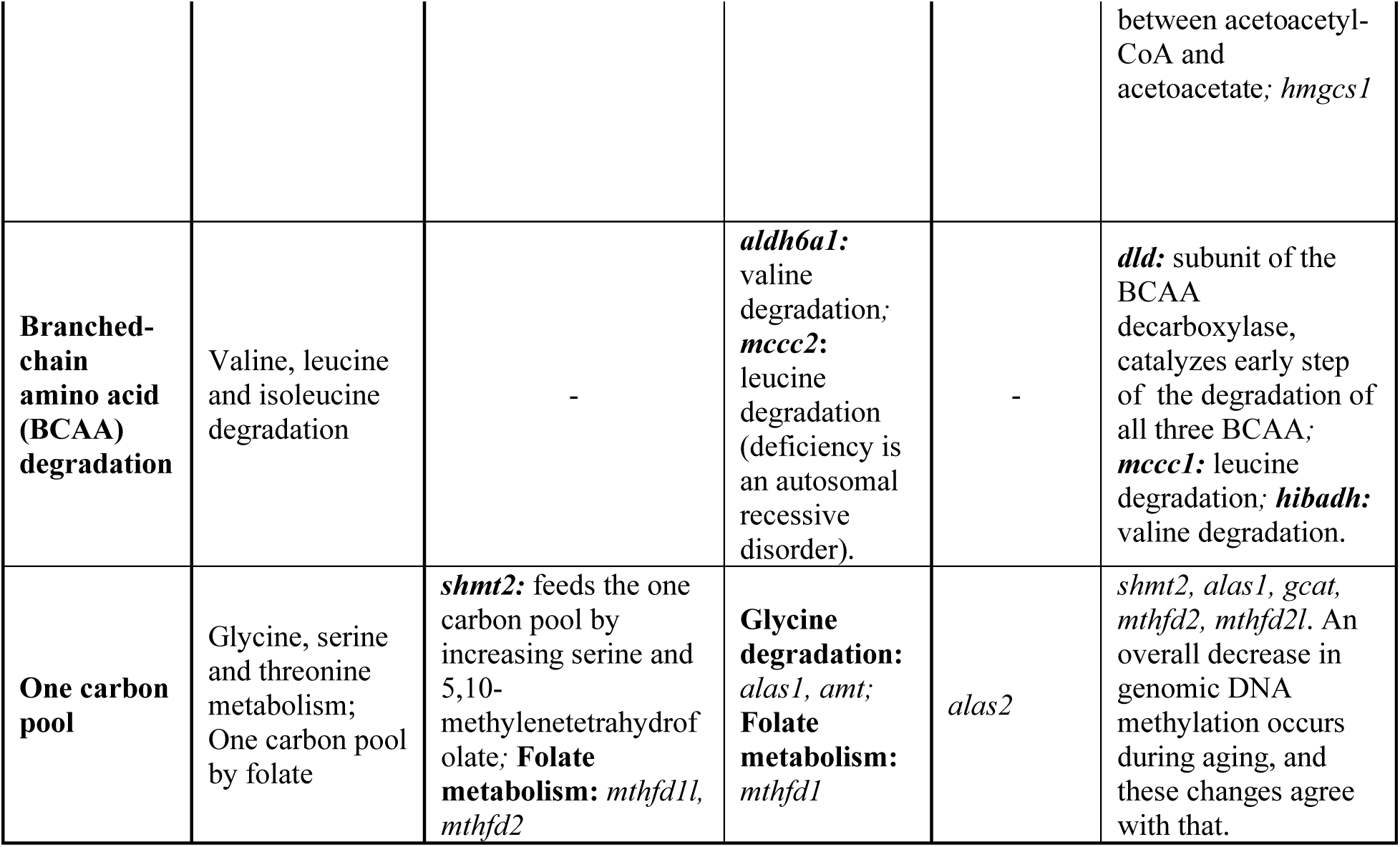
Summary of the main biological processes and pathways identified among differential hub genes during neurotransmission and aging in the neuron.

### Differential hub gene abundance changes in the astrocyte suggest a metabolic switch during brain aging

We next performed the same pathway enrichment of DHG in the astrocyte during neurotransmission and brain aging, followed by manual curation. The number of DHG in the astrocyte was lower than those found in the neuron, leading also to a lower number of enriched pathways. All five biological processes described for the neuron were also found during astrocyte aging.

In the first group we identified DHG enriched in central energy metabolism pathways (Figures 6a and 6a’, blue). These were “Pyruvate metabolism” during both astrocyte neurotransmission and aging, while “Central metabolism in cancer” was only enriched during neurotransmission, and “Citrate cycle (TCA cycle)” was only enriched during astrocyte aging. The following changes were observed in the astrocyte during neurotransmission (Figure 6a). First, *slc2a1*, encoding for the leading glucose uptake transporter in the blood-brain barrier GLUT1 was upregulated. Second, *ldha*, which encodes for a subunit of lactate dehydrogenase (LDH) that favors lactate levels in the interconversion between pyruvate and lactate was also upregulated (Bittar et al., 1996). And third, *pcx* and *acss1*, which encode for enzymes that feed substrates into the TCA cycle, were downregulated. These changes suggest high glucose uptake during neurotransmission by the astrocyte, elevated lactate synthesis, and low TCA cycle flux, which agrees with an active astrocyte-neuron lactate shuttle (ANLS). In contrast, during astrocyte aging, we observed the following changes (Figure 6a’). Instead of *ldha*, we observed upregulation of *ldhb*, which encodes for an LDH subunit that favors pyruvate levels. Furthermore, *mdh1* and *mdh2*, which encode for the cytosolic and mitochondrial malate dehydrogenases, respectively, were upregulated. These enzymes participate in the malate-aspartate shuttle, which transports reducing equivalents into mitochondria (NADH) therefore fueling the electron transport chain and ATP synthesis by oxidative phosphorylation. These changes observed in differential hub gene regulation suggest that oxidative metabolism is favored in the aged astrocyte instead of flux through the ANLS, affecting neuronal energy needs.

**Figure 6.**
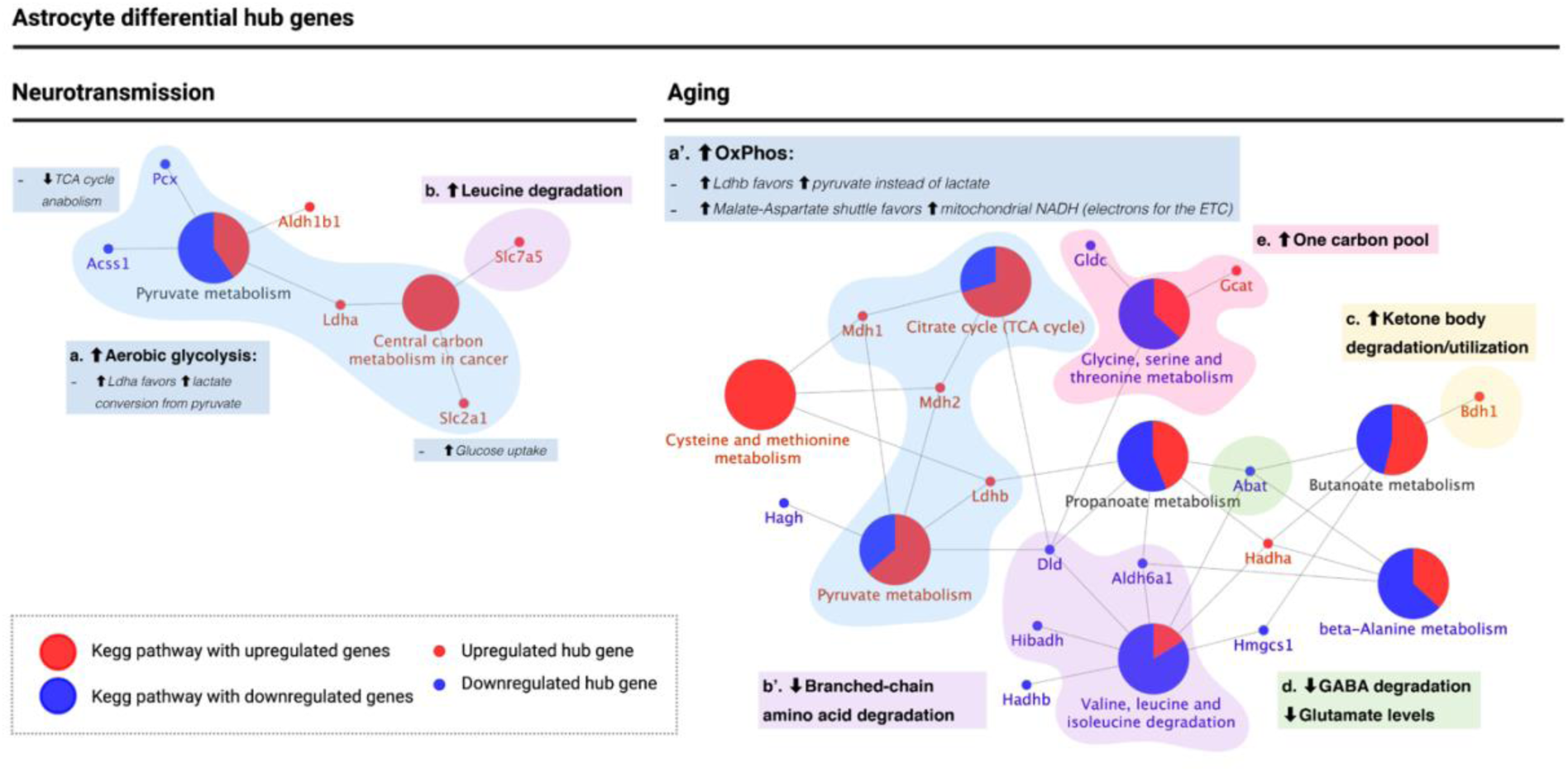
KEGG pathway enrichment analysis of astrocyte differential hub genes suggests a metabolic switch from aerobic glycolysis to oxidative phosphorylation during aging. **a-a’**. Metabolic switch (blue): upregulation of *ldha* during neurotransmission but *ldhb* during aging. *Ldha/b* genes encode for subunits of lactate dehydrogenase, which catalyzes the interconversion of pyruvate into lactate. *Ldha* subunits favor lactate levels and were upregulated during neurotransmission, while *ldhb* favors pyruvate and is upregulated during aging. Also, the major glucose uptake transporter in the blood-brain barrier, encoded by *slc2a1*, was upregulated during neurotransmission only. Instead, during aging, *mdh1/2* encode for enzymes of the malate-aspartate shuttle, which allows transport of NADH into the mitochondrial matrix to provide electrons for the ETC. Both genes were upregulated during aging, in agreement with a high OxPhos rate. **b-b’**. Branched-chain amino acid (BCAA) degradation (purple): during neurotransmission, upregulation of *slc7a5* was observed (involved in leucine degradation), while during aging, three enzymes involved in BCAA degradation, including *dld*, were downregulated. **c**. Ketone body degradation/utilization (yellow): the enzyme encoded by *bdh1* catalyzes the interconversion of acetoacetate and β-hydroxybutyrate, the two main ketone bodies, and was upregulated during aging only. **d**. Synaptic transmission (green): *abat* encodes for an enzyme that breaks down GABA into glutamate and is downregulated during aging in the astrocyte. **e**. One carbon pool (pink): differential hub gene expression associated with one-carbon metabolism suggests an increase in one-carbon pool during astrocyte aging.

The second group was related to branched-chain amino acid (BCAA) degradation (Figures 6b and 6b’, purple), where *slc7a5*, involved in leucine degradation, was upregulated during neurotransmission (Figure 6b, purple). In contrast, *dld*, *aldh6a1,* and *hibadh*, all BCAA degradation genes, were downregulated during brain aging (Figure 6b’, purple). These results are in line with BCAA accumulation in the astrocyte during brain aging.

The third group was associated with ketone body degradation/utilization (Figure 6c, yellow), where *bdh1* was upregulated during brain aging. As mentioned previously, this gene encodes for the enzyme that catalyzes interconversion of acetoacetate and β-hydroxybutyrate, thus suggesting that the aged astrocyte favors ketone body degradation or utilization. The fourth group was associated with glutamate levels (Figure 6d, green), including *abat*, which encodes for an enzyme that degrades GABA converting it into glutamate. *Abat* was found downregulated during astrocyte aging. These results support a metabolic switch in the astrocyte during brain aging that decreases ANLS flux, which is required for meeting the high-energy neuronal demand, promoting ATP synthesis for the astrocyte’s use. Finally, the fifth group was related to one-carbon pool regulation (Figure 6e), where *gcat* was upregulated during aging, while *gldc* was downregulated. *Gcat* activity feeds the one-carbon pool while *gldc* consumes one carbon intermediates. These results strongly suggest that the aging astrocyte favors the one-carbon pool. A summary of all changes observed in the astrocyte is included in Table 2.

**Table 2.** Summary of the main biological processes and pathways identified among differential hub genes during neurotransmission and aging in the astrocyte.

| Process or gene(s) | Associated KEGG pathways and biological processes | Differential hub genes upregulated after neurotransmission | Differential hub genes downregulated after neurotransmission | Differential hub genes upregulated in the aged brain | Differential hub genes downregulated in the aged brain |
| --- | --- | --- | --- | --- | --- |
| **Central energy metabolism: switch from aerobic glycolysis (neurotransmission) into oxidative phosphorylation metabolism** | Pyruvate metabolism; Citrate cycle (TCA cycle); Central carbon metabolism in cancer | **Aerobic glycolysis and astrocyte-neuron lactate shuttle (ANLS):** *ldha*, catalyzes interconversion between pyruvate and lactate, favoring lactate; *slc2a1 (a.k.a. GLUT1)*, the main glucose uptake transporter in the blood brain barrier | *pcx*, *acss1*: feed the TCA cycle with intermediates (TCA anabolic reactions). Also in agreement with aerobic glycolysis metabolic state | **Oxidative metabolism:** *ldhb* favors pyruvate levels instead of lactate (suggesting glycolytic flux in the aged astrocyte is directed towards its own ATP synthesis); *mdh1* and *mdh2* are involved in the malate-aspartate shuttle, which allows NADH transport into the mitochondria to provide electrons for the electron transport chain (also in agreement with favoring ATP synthesis in the aged astrocyte instead of the ANLS). | - |
| **Synaptic activity and glutamate** | Na/K-ATPase pumps; glutamate synthesis | - | - | - | *abat*: catalyzes GABA degradation, producing glutamate. |
| **Ketone body metabolism** | Synthesis and degradation of ketone bodies | - | - | *bdh1*: catalyzes interconversion of the two main ketone bodies, acetoacetate and beta-hydroxybutyrate (this conversion is required for ketone body utilization). | - |
| **Branched-chain amino acid (BCAA) degradation** | Valine, leucine and isoleucine degradation | *slc7a5* is involved in leucine (a BCAA) degradation | - | - | *dld*: subunit of the BCAA decarboxylase, catalyzes early step of the degradation of all three BCAA; *aldh6a1* and *hibadh*: valine degradation |
| **One carbon pool levels** | Glycine, serine and threonine metabolism; One carbon pool by folate | - | - | *gcat*: threonine degradation into glycine and acetyl-CoA | *gldc*: glycine degradation |

## DISCUSSION

In the present work, we analyzed the neuron-astrocyte metabolic network by integrating a flux-based approach (Flux Balance Analysis), a centrality analysis, which addresses the intrinsic structure of the network. This network analysis was followed by cross-reference of the identified *hub genes*, with gene expression data for both cell types during neurotransmission and brain aging (see *Workflow Overview* in the Results section, and Figures 1 and 2). The integration of these three approaches allowed the identification of *differential hub genes (DHG)*, which are a robust selection of gene candidates with a high probability of playing pivotal roles in the neuron-astrocyte metabolic network, and to explain the molecular mechanisms of age-associated brain functional decline. DHG were further analyzed using pathway enrichment analysis. This allowed identifying the main biological processes in which DHG participate during neurotransmission and/or brain aging.

### Impaired central energy metabolism in the aged neuron

Brain energy metabolism dysfunction has been described as a hallmark of brain aging (Cunnane et al., 2020; Mattson and Arumugam, 2018), and metabolic deficit in the neuron during human brain aging has been reported (Boumezbeur et al., 2010). This group reported that flux through the tricarboxylic acid (TCA) cycle decreased by 28% in the presynaptic neuron using *in vivo* magnetic resonance spectroscopy. However, the genes involved in this deficit remain largely unknown. Our analyses showed that the aging neuron downregulated a high number of TCA cycle genes, including:

1. Subunits of the pyruvate dehydrogenase complex, *dld* (EC:1.8.1.4)*, pdha1* (EC:1.2.4.1)*, pdhb* (EC:1.2.4.1), and *dlat* (EC:2.3.1.12), which catalyzes the conversion of pyruvate into acetyl-CoA for entry into the TCA cycle. Among these, it is worth highlighting that *dld* (EC:1.8.1.4) is also a catalytic subunit of two other essential dehydrogenase complexes: the α-ketoglutarate dehydrogenase complex (α-KGDH), which catalyzes the conversion from α-KG into succinyl-CoA (reaction that produces NADH in the mitochondrial matrix), and the branched-chain amino acid (BCAA) dehydrogenase complex. Remarkably, downregulation of *dld* severely affects overall metabolic function and causes the hereditary disease dihydrolipoamide dehydrogenase deficiency (OMIM: 246900) (Brautigam et al., 2006; Odievre et al., 2005). However, the role of *dld* during brain aging has not been described.
2. Two isoforms of malate dehydrogenase (*mdh1* and *mdh2*) were downregulated in aging neurons. These enzymes are involved in the malate-aspartate shuttle (MAS), which allows the shuttling of NADH into the mitochondrial matrix (Minarik et al., 2002), providing reducing equivalents for the electron transport chain (ETC). Furthermore, it has been shown that the expression of malate-aspartate shuttle enzymes decreases with normal aging and can be reverted using dietary restriction (Goyary and Sharma, 2008). Also, *loss-of-function* mutations in the *mdh2* gene are associated with severe neurological deficits in children (Ait-El-Mkadem et al., 2017). Notably, the NADH/NAD+ ratio is one of the driving forces of the ANLS together with pyruvate levels (Weber and Barros, 2015), highlighting these two enzymes as candidates to study age-associated brain functional decline.
3. Downregulation of *idh1* and *idh2* (encoding for the enzymes Isocitrate Dehydrogenases 1 and 2), which catalyze α-ketoglutarate (α-KG) synthesis. This metabolite exits the TCA cycle and is converted into glutamate, which is the main excitatory neurotransmitter, and therefore it is central for metabolism since it connects energy metabolism with neurotransmission via glutamate.

These changes agree with previous findings of metabolic deficit in the neuron during aging and provide both previously reported genes (validating our modeling method) and novel gene targets. An energetic shortage in a cell with such high energy demand is critical and will necessarily lead to dysfunction.

### Astrocyte metabolic switch from aerobic glycolysis to oxidative phosphorylation

Differential gene expression patterns in astrocytes indicate a metabolic switch from aerobic glycolysis to oxidative metabolism. Since astrocytes fuel neurons with lactate, this metabolic switch can lead to neuronal energy deficit. This behavior has been described previously as a *selfish phenotype* adopted by the astrocyte during aging (Jiang and Cadenas, 2014). Here, astrocytes use pyruvate for their ATP synthesis instead of shuttling it to neurons. In this regard, lactate dehydrogenase isoforms dictate the fate of pyruvate, either by favoring its conversion into lactate or by directing it into the TCA. Specifically, lactate dehydrogenase isoform LDH-5 favors lactate production, while isoform LDH-1 favors pyruvate production (Bittar *et al.*, 1996). Consistent with the ANLS, the *ldha1* gene (coding for polypeptides forming the LDH-5) increases in astrocytes during neurotransmission. Remarkably, the *ldhb* gene, which codes for the subunits of the polypeptides of LDH-1, was upregulated during brain aging. These findings support the glycolytic-to-oxidative metabolic switch in the astrocyte. Furthermore, while LDH-1 isoenzymes localize in both neurons and astrocytes, the LDH-5 isoenzymes localize exclusively in astrocytes (Bittar *et al.*, 1996). Hence, it is highly relevant that *ldhb* increases in aged astrocytes.

*Ldh* upregulation occurs *Drosophila melanogaster* aging, where *loss-of-function* in either neurons or astrocytes leads to an increase in lifespan, while *gain-of-function* reduces lifespan (Long et al., 2020). *Ldh* overexpression also leads to increased neurodegeneration and motor function decline, while downregulation is neuroprotective (Long *et al.*, 2020). However, specific studies on the *ldha-to-ldhb* switch in astrocytes have not been performed and would be of great interest to understand the mechanisms of functional brain decline during aging.

Returning to MAS, *mdh1* and *mdh2* were upregulated during astrocyte aging (as opposed to downregulation in the aged neuron), supporting the oxidative metabolism switch in the astrocyte (Minarik *et al.*, 2002). Of note, while there were controversial reports of Aralar, the glutamate/aspartate antiporter in the MAS not being expressed in astrocytes (McKenna et al., 2006), later evidence showed the opposite (Li et al., 2012). In fact, both transcriptomic databases used here detected *slc25a12* transcript expression (which encodes for Aralar) in astrocytes, albeit not differentially expressed (Consortium, 2020; Hasel *et al.*, 2017).

Taken together, these results show that while the neuron displays an intrinsic energetic deficit as demonstrated by its expression changes, the astrocyte further contributes to this deficit by undergoing a metabolic switch into a *selfish phenotype* during brain aging.

### Role of mdh2 and ldhb in the metabolic switch of other cell types

In cancer cells, the “Warburg effect”, which is also known as aerobic glycolysis, was first described. In the transition from normal-to-tumoral cells, they undergo an oxidative-to-aerobic glycolysis switch, favoring proliferation (Vander Heiden et al., 2009). However, exposure of cancer cells to radiation induces a switch to oxidative metabolism arresting proliferation (Lu et al., 2015). Notably, treatment of cancer cells with an Mdh2 inhibitor induces downregulation of oxidative phosphorylation (Ban et al., 2016), which is in line with the role in the metabolic switch of Mdh2.

Furthermore, it was recently reported that *ldhb* plays a role in tumor-associated macrophages in breast carcinoma (Frank et al., 2021). These macrophages express low levels of *ldhb*, perform aerobic glycolysis and secrete high lactate levels. Yet, when the authors upregulate *ldhb* this significantly decreases lactate production in these macrophages, further supporting the role of *ldhb* upregulation in inducing an oxidative phenotype.

### Impaired branched-chain amino acid degradation

Valine, leucine, and isoleucine are the three branched-chain amino acids (BCAA). Impairment in their degradation is detrimental to overall metabolic health (Richardson et al., 2021; Solon-Biet et al., 2019; Yu et al., 2021). We observed the downregulation of genes involved in BCAA degradation during neurotransmission and aging in the neuron. During neurotransmission, *aldh6a1*, involved in valine degradation (Figure 5e and Table 1), and *mccc2* involved in leucine degradation, while during aging, *mccc1* (also leucine degradation), *hibadh* (valine degradation), and *dld*, which catalyzes an early step in the degradation of all three BCAA (Figure 3e’). Importantly, a recent report showed that detrimental effects of BCAA are mediated mainly through isoleucine, and to a lesser extent, by valine (Yu *et al.*, 2021).

Significantly, *dld* was also downregulated in the astrocyte. In fact, *slc7a5* involved in leucine degradation was upregulated during neurotransmission in the astrocyte, while *dld*, *aldh6a1,* and *hibadh* were downregulated in the aging astrocyte. Taken together with the fact that *dld* encodes for a subunit in three central dehydrogenases, its downregulation in both the aged neuron and the astrocyte, plus its role in BCAA degradation, we propose *dld* as one of the strongest candidates to target in the aging brain.

### Altered ketone body metabolism

Ketone bodies are produced during caloric restriction, which is the only intervention known to extend lifespan across various organisms (Mattison et al., 2017), and several metabolic challenges are being developed to emulate the effects of caloric restriction, including the ketogenic diet (Newman et al., 2017; Roberts et al., 2017) and intermittent fasting (Dias et al., 2021). Our results suggest that DHGs are regulated in the neuron such that during neurotransmission, they suggest downregulation of ketogenesis by upregulation of *hmgcs1*, the cytosolic isoform of *hmgcs2*. While Hmgcs2 (the mitochondrial isoform) catalyzes the first rate-limiting step in ketogenesis (Hegardt, 1999), Hmgcs1 catalyzes cholesterol biosynthesis in the cytosol instead of ketone body synthesis. However, during neuronal aging, *hmgcs2* was upregulated while *hmgcs1* downregulated, thus suggesting upregulation of ketone body synthesis (Hegardt, 1999). *Bdh1* (EC:1.1.1.30) and *oxct1* (EC:2.8.3.5), which participate in the utilization of ketone bodies were downregulated, suggesting downregulation of ketone body degradation during neuronal aging.

In the astrocyte, *bdh1* (EC:1.1.1.30) was upregulated. This gene encodes for the enzyme that catalyzes the interconversion between β-hydroxybutyrate and acetoacetate, the two main ketone bodies, a reaction required for acetoacetate conversion into acetyl-CoA for eventual ATP synthesis (Cunnane *et al.*, 2020). Therefore, this suggests an upregulation of ketone body degradation and utilization in the astrocyte during aging. Taken together, our results suggest that ketone body utilization increased during astrocyte aging while the aging neuron upregulated ketogenesis. These results agree with those mentioned above regarding astrocyte energy expenditure being favored over the neuronal demand during brain aging.

### Downregulation of genes associated with synaptic transmission in the aging neuron

During synaptic transmission, we observed that the neuron upregulated genes encoding for four sodium/potassium-ATPase (Na/K-ATPase) pumps (Figure 5b, orange), while four out of five Na/K-ATPase pumps were downregulated during neuronal aging (Figure 5b’, orange). These pumps are required to re-establish neuronal ion gradients after neurotransmission (Baeza-Lehnert *et al.*, 2018; Erecinska and Silver, 1994). A downregulation of their expression during aging could contribute to neuronal dysfunction. Furthermore, *got1* (EC:2.6.1.1), which synthesizes glutamate from TCA intermediate α-ketoglutarate, is upregulated during neurotransmission, while *got1* and *got2* (EC:2.6.1.1) were downregulated during brain aging. Since glutamate is the main excitatory neurotransmitter (Reiner and Levitz, 2018) and α-ketoglutarate a key metabolic intermediate in the TCA cycle, these two enzymes, in particular, *got1* (the cytoplasmic isozyme), with opposite regulation during neurotransmission and aging, provide a link between central energy metabolism and synaptic activity. Remarkably, activity for the enzyme encoded by *got* is increased in the brain of Alzheimer’s disease individuals compared with healthy controls (D’Aniello et al., 2005). However, further characterization of the enzyme during pathological or healthy brain aging is still lacking, making it an exciting target for future studies.

Regarding glutamate levels, the enzyme *abat* (EC:2.6.1.19), which also catalyzes the conversion of α-ketoglutarate into glutamate, is downregulated during astrocyte aging. The reaction catalyzed by this enzyme involves the degradation of ɣ-aminobutyric acid, or GABA, the main inhibitory neurotransmitter.

As a whole, glutamate synthesized by *got1*, *got2,* and *abat* decreases during both neuron and astrocyte aging. This has a possible detrimental effect on neurotransmission and coupled with the downregulation of the expression of Na/K-ATPase pumps in the aging neuron, provides valuable future horizons for elucidating the molecular mechanisms of brain aging.

### Altered one-carbon pool for tetrahydrofolate (THF) synthesis

The final group we observed was defined by KEGG pathways “One carbon pool by folate” (KEGG map00670) and “Glycine, serine and threonine metabolism” (KEGG map00260). These pathways are of interest during brain aging because THF is the precursor for S-adenosylmethionine (SAM), the substrate required for methylation, including DNA and histone methylation (Jones et al., 2015; McCauley and Dang, 2014; Sen et al., 2016) linking central energy metabolism with epigenetic modifications. Overall methylation levels decrease during aging (Benayoun et al., 2015; Levine et al., 2020; Wang et al., 2020) and is one of the epigenetic clocks, which can be modified by metabolic challenges such as caloric restriction (Gensous et al., 2019). Furthermore, glycine and serine degradation also feed the one-carbon pool (Amelio et al., 2014).

In the neuron, we observed the following changes in one carbon pool associated enzymes (Figures 5f and 5f’). During neurotransmission, *shmt2* (EC:2.1.2.1), an enzyme that synthesizes 5,10-Methylene-THF (KEGG map00260), *mthfd1l* (EC: 6.3.4.3) and *mthfd2* (EC: 3.5.4.9) were all upregulated. The enzymes encoded by *mthd1l* and *mthfd2* feed the THF pool (KEGG map00670). However, in the aged neuron, *alas1* was downregulated and this change is associated with decreased THF levels (Amelio *et al.*, 2014). These changes suggest that during neurotransmission, availability of THF increases, while during aging it decreases, in line with the overall decrease in DNA methylation reported during aging (Benayoun *et al.*, 2015; Levine *et al.*, 2020; Wang *et al.*, 2020; Zampieri et al., 2015). Furthermore, THF is required for glutathione synthesis (GSH), required to quench the high reactive oxygen species levels produced from oxidative phosphorylation. Therefore, high THF levels can be associated with the oxidative metabolism during neurotransmission.

In the astrocyte, we observed differential expression of *gcat* and *gldc*. The Gcat enzyme (EC:2.3.1.29) increases glycine levels from threonine degradation, and therefore feeds the one carbon and THF pool was upregulated during aging. In contrast, Gldc (EC:1.4.4.2) catalyzes glycine degradation and therefore consumes THF, was downregulated. These results suggest an overall increase in the one carbon pool during astrocyte aging, which also suggest an increase in THF and a subsequent increase in availability for GSH synthesis. This agrees with an oxidative metabolic state that is also in line with the aerobic glycolysis to oxidative phosphorylation switch we propose for the aged astrocyte.

### Labor division between the neuron and astrocyte

Differential hub gene expression in the aged astrocyte further reinforces the notion of division of labor between the neuron and astrocyte in the metabolic network shown by both flux balance analysis and centrality analysis. The regulation of biological processes associated with DHG suggests that the aged astrocyte fails to perform its part in this division of labor, which is mainly providing lactate to the neuron and recycling glutamate and glutamine. Instead, the astrocyte switches into a *selfish phenotype*, where energy expenditure reallocates to this cell during brain aging. Taken together, differential hub gene regulation in both cell types strongly supports neuronal metabolic deficit, which could contribute to the cognitive deficit observed in the brain during aging.

## CONCLUDING REMARKS

The work reported here employed two network-based approaches combined with bioinformatics transcriptomics analyses. All numerical predictions were cross validated using reported experimental data. The strategy we developed will be of use for identifying hub genes pivotal for the function of other cellular systems. Our findings suggest that the astrocyte undergoes a metabolic switch from aerobic glycolysis to oxidative metabolism, with a concomitant upregulation of THF precursor synthesis required for glutathione synthesis to control the increased oxidative stress caused by this metabolic switch. Additionally, DHG in the neuron show massive metabolic impairment and downregulation of genes required for synaptic transmission. The genes and pathways identified here include genes previously associated with brain aging and novel targets for aging prevention.

## ACKNOWLEDGEMENTS

This work was funded by: FONDAP Geroscience Center for Brain Health and Metabolism (15150012 to C.G.-B.); ANID FONDECYT Postdoctoral 3180180 (to D.L.-L.) and 3200726 (to A.A.); ANID FONDECYT Regular 1200029 (to L.F.B.); ANID FONDECYT Regular 1211902 and CEDENNA through Financiamiento Basal para Centros Científicos y Tecnológicos de Excelencia (grant AFB180001) (to F.T. and M.K.); and Beca Santander Movilidad Internacional Profesores (to F.T.).

We also thank Manuel I. Muñoz-González for developing and implementing the code for high-performance computing simulations. We thank the National Laboratory for High-Performance Computing (NLHPC) of Chile; Powered@NLHPC: This research was partially supported by the supercomputing infrastructure of the NLHPC (ECM-02).

## AUTHOR CONTRIBUTIONS

A.A. and D.L.L. conceived, designed the study, performed data analysis and interpretation, prepared figures and wrote the manuscript. A.A. performed flux and centrality analyses. D.L.-L. performed transcriptomic and functional analyses. F.T. and M.K. contributed to the design and computation of centrality analysis. F.B.-L and L.F.B. contributed empirical metabolite measurements and their interpretation. C.G.-B. contributed to study conceptualization and data interpretation. All authors discussed the results and critically revised and edited the manuscript.

## METHODS

### Modeling rationale

Our modeling approach tackled three aspects of the metabolic network conformed by neurons and astrocytes: (i) fast response to glutamatergic-neurotransmission workload (Ruminot et al., 2019), (ii) constant energy availability, i.e., invariant neuronal concentrations of cytosolic ATP and ADP (Baeza-Lehnert *et al.*, 2018), and (iii) long-term impairment upon aging (Mattson and Arumugam, 2018). We addressed the former aspect (i) by employing a genome-scale constraint-based model of the neuron-astrocyte metabolic network (Lewis *et al.*, 2010); henceforth, the neuron-astrocyte model. To simulate the response to neurotransmission workload (i), we coupled and maximized three critical fluxes. These fluxes were those that are activated under glutamatergic neurotransmission and comprised neuronal ATP consumption derived from sodium removal, the ANLS, and the GGC. These three events were combined into a single flux denoted as the metabolic objective. The second aspect (ii), constant energy availability, was managed by subjecting the maximization of the metabolic objective to a steady-state constraint. This optimization-based procedure is known as FBA (Orth et al., 2010) and simulates an optimal metabolic response to neurotransmission. The essentials of FBA can be found in the Supplemental Theoretical Framework Section 1. The FBA allowed us to identify the optimal metabolic reactions, which were the reactions responsible for achieving a proper response to neurotransmission. Up to this point, our model encoded the stationary and optimal nature of the response to neurotransmission workload. Notably, metabolic states computed via FBA simulate events that are required to be reproducible for the cell (Fell, 2021). Consistently, the brain must maintain a reproducible outcome, namely a proper response to energy workload, particularly in the face of aging. Even though part of aging-derived damage to brain metabolism may reside in fast stationary events, much of aging deterioration may relate to non-stationary long-term events. Network topology can encode wide-spectrum phenomena beyond steady-state and short timescales since it can encode the row space of the stoichiometric matrix (see Supplemental Theoretical Framework Section 4). Therefore, we identified a group of reactions that modulate the optimal metabolic response via topological effects to analyze aging-derived phenomena (theoretical details on the topology-based analysis are exposed in the Supplemental Theoretical Framework section 2). Since we employed centrality analysis (Borgatti, 2006), these modulators were called central metabolic reactions and, along with the optimal metabolic reactions, were used to identify aging-affected genes.

### Neuron-astrocyte metabolic network

We used a genome-scale metabolic network reconstruction (Fang et al., 2020; Thiele and Palsson, 2010) of the glutamatergic synapse comprising neurons and astrocytes (Lewis *et al.*, 2010). This model is available at https://systemsbiology.ucsd.edu/InSilicoOrganisms/Brain.

### Flux constraints

The theory behind constraint-based modeling and flux constraints is briefly presented in the Supplemental Theoretical Framework sections 1.1 to 1.4. Neuronal flux constraints were derived from measurements taken in primary cultures reported by (Baeza-Lehnert *et al.*, 2018). In this study, they used genetically encoded fluorescence resonance energy transfer (FRET) reporters (San Martin et al., 2014; San Martin et al., 2013) along with ion-sensitive dyes to make real-time measurements of intracellular fluxes in neurons co-cultured with astrocytes. (Baeza-Lehnert *et al.*, 2018) investigated how the neuronal ATP pool is maintained upon acute energy demands derived from the activity of the Na+/K+ ATPase pump induced by neuronal stimulation. They were able to estimate that sodium ions are extruded at a rate of 350 µM/s after neuronal stimulation. This sodium efflux rate corresponds to 116.6 µM/s of ATP consumption since 1 molecule of ATP is spent to export 3 ions of sodium. Also, (Baeza-Lehnert *et al.*, 2018) estimated a housekeeping ATP demand of 38 µM/s. Adding the ATP spent during stimuli-associated sodium removal and the housekeeping demand, (Baeza-Lehnert *et al.*, 2018) estimated a total ATP demand of 155 µM/s to re-establish ions gradient after neurotransmision. Also, they reported that at resting conditions neuronal glucose consumption was near 0.9 µM/s (in the presence of lactate) and that neuronal glycolytic rate increases 2.353 times after stimulation. This yields a glycolytic rate of 2.1177 µM/s in stimulated neurons. Overall, a stimulated neuron must cope with an ATP demand of 155 µM/s having a glycolytic rate of 2.1177 µM/s. Considering this glycolytic rate of 2.1177 µM/s and an energy yield of 31 molecules of ATP per glucose (Baeza-Lehnert et al., 2018), the neuronal metabolism roughly produces 66 µM/s of ATP. The rest of the required ATP is achieved via lactate uptake, where lactate is supplied by astrocytes (Magistretti and Allaman, 2018). Such lactate production in astrocytes is associated with an astrocytic glycolic flux that is triggered under neuronal stimulation (Fernández-Moncada *et al.*, 2018). We used Flux Balance Analysis (FBA) to fit the astrocytic glycolytic flux to meet the neuronal ATP demand of 155 µM/s, and hence the sodium efflux of 350 µM/s. Astrocytic oxygen uptake was fixed at 0.01666 µM/s as reported in experiments where astrocytes are co-cultured with neurons that undergo stimulation (Fernández-Moncada *et al.*, 2018). Hence, we computed the optimal metabolic state using the latter astrocytic oxygen uptake rate along with the fitted astrocytic glycolytic rate, the neuronal glycolytic rate, and the housekeeping ATP demand as flux constraints.

### Phenotypic phase plane analysis

In this analysis, a non-zero slope of the planes means that the optimal state depends on the given substrates (Edwards et al., 2002). We computed the phenotypic phase planes as follows:

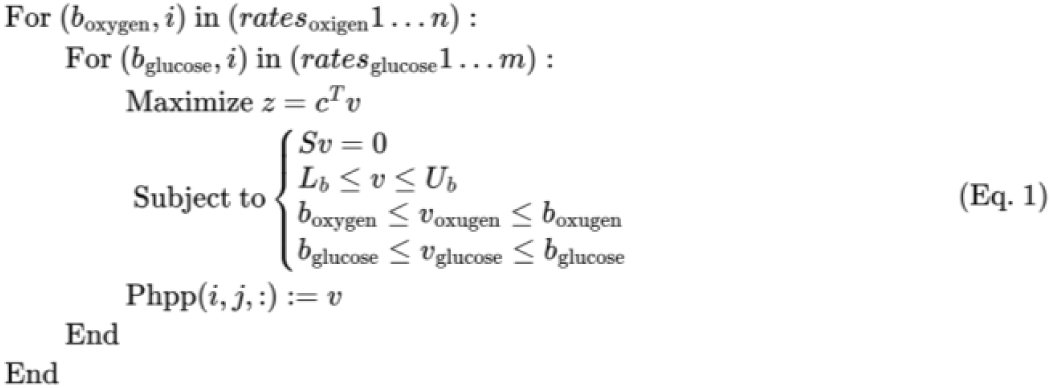

here, *rates_oxygen_* and *rates_glucose_* are ranges of uptake rates which may be of any length. For details, the theoretical basics of the Flux Balance Analysis (FBA) are presented in the Supplemental Theoretical Framework sections 1.2 to 1.4. The term *z* = *C^T^v* correspond to the metabolic objective of the FBA, which correspond to a linear combination of the fluxes *V* weighted by *C^T^*. Specifically, the vector *C* has ones for the reactions shown in Figures 2a-c, and zero for the rest. The stoichiometric matrix is denoted as *S*, and the equality constraint *SV* = 0 corresponds to the mass-balance at steady state. The vectors *L_b_*, *U_b_*, *b_oxygen_*, *b_glucose_* are bounds for the inequality constraints, respectively, these correspond to the full-length lower bounds, full-length upper bounds, lower bound for oxygen uptake rate, and lower bound for glucose uptake rate. The flux variables *V_oxygen_* and *V_glucose_* correspond to oxygen uptake rate, and glucose uptake rate. The term *Phpp* is a tensor where the first two dimensions are of the corresponding lengths of the uptake ranges. The third dimension of the tensor *Phpp* is the number of fluxes in the model. From this tensor, we extracted the phenotypic phase planes shown in Figures 2d -g.

### Sensitivity analysis of the FBA

The sensitivity analysis was carried out over the solution of the FBA. This was done via calculation of what is known as the reduced cost vector (Maarleveld et al., 2013; Price et al., 2004). Each value of this vector indicates the amount by which the objective function changes upon an increase in a given flux. Thus, a reduced cost (*δ_i_*) is the sensitivity of the objective function *z* with respect to a change in the *i*th flux value (*v_i_*):

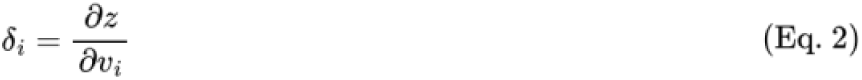

Hence, a group of reactions able to perturb the optimal response may be identified. This group comprised of reactions having non-zero *δ_i_*, being named as the *sensitivity set*. This group acts as an "interface" able to send fast perturbations to the optimal state.

### Absolute Optimality

We constructed an index of reaction importance in the context of the optimal metabolic response. This index was called Absolute Optimality (AO) and corresponds to the L2 norm of a vector composed by normalized flux and normalized sensitivity. We normalized flux and sensitivity in order to get standardized positive values. Such a normalization consisted in appling the Signed Pseudo Logarithm and rescaling the values to a zero-one range (*scaler*):

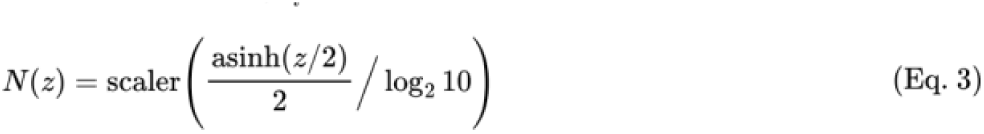

where *z* corresponds to any given flux or sensitivity. Then, the AO for the ***i*** reaction is:

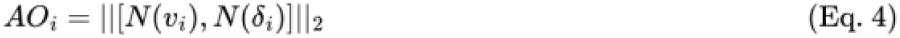

where *v_i_*, and *δ_i_* corresponds to the flux and sensitivity of the ***i*** reaction, respectively.

### Absolute Centrality Contribution

We carried out centrality analysis over the reaction projection of the stoichiometric matrix. The projection of the stoichiometric matrix is explained in detail in the Supplemental Theoretical Framework section 2.2. It is worth noting that we did not only assess the centrality of the reactions involved in the sensitivity set. Rather, we determined how other nodes contribute to the centrality of the reactions involved in the sensitivity set. In this sense, we build from the concept of *induced centrality,* which views a node’s centrality as a measure of its contribution to another node’s centrality (Borgatti, 2010). Formally, induced centrality accounts for the contribution of any node to the network’s cohesiveness, where cohesiveness is defined as the aggregation of all nodes’ centrality scores. Induced centrality is computed by taking any centrality metric and aggregating all node scores (averaging them, for instance) to get a baseline measure of network cohesiveness, and then recalculating the aggregation without the node of interest. The difference between the baseline and the recalculation yields the induced centrality of the node of interest. We adapted this procedure to our ends. Instead of taking the centrality scores of all nodes, we only took sensitivity nodes and aggregated them via arithmetic mean. Also, instead of using the difference, we used the fold change between the baseline and the recalculation. Formally, our implementation of the node induced centrality defines the basal centrality of the sensitivity set as the mean of its node centralities:

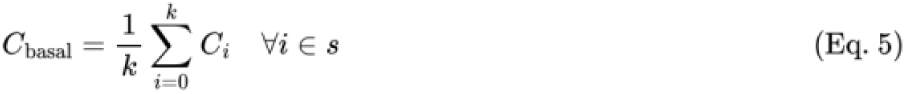

Where *C* is any given centrality metric (eigenvector, closeness or information), and *k* is the numbers of members of the the sensitivity set (*s*), while *C_i_* is the centrality of a member of the sensitivity set. Next, we defined the perturbed centrality of the sensitivity set as the same mean but recalculated without node *x*,

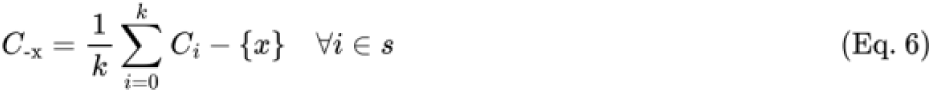

here, *C_i_* − {*x*} refers to the recalculated centrality (centrality without *x*) of a member of sensitivity set, and *C_−x_* is the perturbed centrality. Then, we computed the induced centrality of node *x* as,

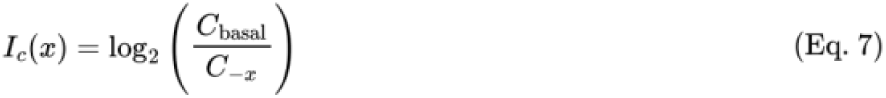

where centrality *C* can be eigenvector, closeness or information centrality. *I_c_*(*x*) is the contribution of node *x* to the centrality of sensitivity set. Next, we normalized these data in order to get standardized positive values. To such an end, we employed Eq. 3. Induced centrality may be calculated by using centrality metrics that inform on the probability of getting an interaction (eigenvector) or calculated via centralities associated with the cost of such interaction (closeness or information centrality). Details on the concept of probability and cost-associated centralities can be found in the Supplemental Theoretical section 3. Finally, we added the normalized cost-associated induced centralities into one quantity (*C_S_*(*x*)), *P_S_*(*x*)

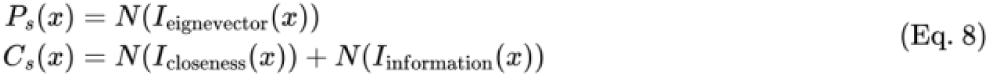

here, subscripts *s* highlight the fact that induced centralities are defined regarding the sensitivity set. Finally, we computed the Absolute Centrality Contribution (ACC) as the L2 norm of a vector compose by the probability and the cost,

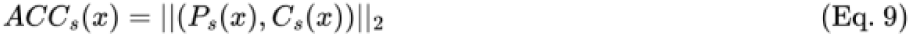

here, *ACC_s_*(*x*) encodes the contribution of node *x* to the availability of the sensitivity set (*s*) to have interactions with the rest of the network.

### Pairwise correlations between nodal contributions

Correlations were calculated using Pearson’s coefficient via its implementation in the R language. Non-parametric coefficients were not necessary as each node was related only to four data points, each one corresponding a different *I_c_*(*x*).

### Hierarchical clustering

We used unsupervised hierarchical clustering to verify the opposite regulation found between neurons and astrocytes regarding their induced centralities. In this sense, we determined if the clusterization of nodal contributions resembles the two-cell structure (neuron-astrocyte) of the network. To this end, each reaction was regarded as a variable while its four induced centralities were regarded as samples. Hence, we computed the correlation matrix between reactions. If there is opposite regulation, the neuron-astrocyte structure should emerge from unsupervised clusterization of the correlation matrix. Hierarchical clustering was done by using euclidean norm to compute distances, and complete-linkage as agglomeration method. The PCA was applied according to standard implementation.

### Genes associated with reactions

Each reaction (enzyme or transporter) is associated with some gene or group of genes. We manually annotated those genes by using the Virtual Metabolic Human website (https://www.vmh.life), which is a database based on information provided by constraint-based stoichiometric models of human metabolism (Noronha et al., 2019).

### Software, programming languages and libraries

Pathway visualizations shown in Figures 2a-c were done using *Escher* (https://escher.github.io/). Phenotypic phase planes (PhPPs) were computed in the Python language using CobraPy (https://opencobra.github.io/cobrapy/). Al l statistical tests (wilcoxon) were carried out employing the ggplot2 built-in function stat_compare_means. All plots shown in Figure 2 and 3 were composed and rendered using the R language (https://www.r-project.org/) employing the library ggplot2 (https://ggplot2.tidyverse.org/), except for the network visualizations shown in Figures 2h, i (left-side), and j (left-side) which were made in Python using graph-tool (https://graph-tool.skewed.de/). Hierarchical clustering and heatmap were done using the R library ComplexHeatmap (https://jokergoo.github.io/ComplexHeatmap-reference/). In the same manner, PCA was carried out in R by using the library PCAtools (https://github.com/kevinblighe/PCAtools).

### Code availability

The code to replicate the results presented in Figures 3 and 4 is available under prior solicitation to the corresponding author.

### High-performance computing software and infrastructure

This research was partially supported by the supercomputing infrastructure of the National Laboratory for High Performance Computing (NLHPC) of Chile (ECM-02). Distributed computing was implemented by using Python package Ray (https://docs.ray.io/).here, *ACC_s_*(*x*) encodes the contribution of node *x* to the availability of the sensitivity set (*s*) to have interactions with the rest of the network.

### Pairwise correlations between nodal contributions

Correlations were calculated using Pearson’s coefficient via its implementation in the R language. Non-parametric coefficients were not necessary as each node was related only to four data points, each one corresponding a different *I_c_*(*x*).

### Hierarchical clustering

We used unsupervised hierarchical clustering to verify the opposite regulation found between neurons and astrocytes regarding their induced centralities. In this sense, we determined if the clusterization of nodal contributions resembles the two-cell structure (neuron-astrocyte) of the network. To this end, each reaction was regarded as a variable while its four induced centralities were regarded as samples. Hence, we computed the correlation matrix between reactions. If there is opposite regulation, the neuron-astrocyte structure should emerge from unsupervised clusterization of the correlation matrix. Hierarchical clustering was done by using euclidean norm to compute distances, and complete-linkage as agglomeration method. The PCA was applied according to standard implementation.

### Software, programming languages and libraries

Pathway visualizations shown in Figures 2a-c were done using *Escher* (https://escher.github.io/). Phenotypic phase planes (PhPPs) were computed in the Python language using CobraPy (https://opencobra.github.io/cobrapy/). All statistical tests (wilcoxon) were carried out employing the ggplot2 built-in function stat_compare_means. All plots shown in Figure 2 and 3 were composed and rendered using the R language (https://www.r-project.org/) employing the library ggplot2 (https://ggplot2.tidyverse.org/), except for the network visualizations shown in Figures 2h, i (left-side), and j (left-side) which were made in Python using graph-tool (https://graph-tool.skewed.de/). Hierarchical clustering and heatmap were done using the R library ComplexHeatmap (https://jokergoo.github.io/ComplexHeatmap-reference/). In the same manner, PCA was carried out in R by using the library PCAtools (https://github.com/kevinblighe/PCAtools).

### Code availability

The code to replicate the results presented in Figures 3 and 4 is available under prior solicitation to the corresponding author Dasfne Lee-Liu.

### High-performance computing software and infrastructure

This research was partially supported by the supercomputing infrastructure of the National Laboratory for High Performance Computing (NLHPC) of Chile (ECM-02). Distributed computing was implemented by using Python package Ray (https://docs.ray.io/).

### Extracting differential gene expression values from databases

Genes displaying differential abundance in response to glutamatergic neurotransmission were extracted from Supplemental Material reported in (Hasel *et al.*, 2017), using the following threshold reported for the astrocyte: fold-change (stimulated/basal) ≥1.3 or ≤0.77 and p-adjusted-SSS-value < 0.05. Differentially abundant genes reported with or without TBOA treatment were merged into a single gene set. For the neuron, the same parameters were used for comparable results. The same procedure was used to extract genes showing differential abundance in response to brain aging in the astrocyte and neuron (Consortium, 2020). We used the threshold reported by the authors at: age coefficient threshold at 0.005 reported by authors as equivalent to a 10%-fold change and an FDR cutoff of 0.01. Given that the abovementioned studies used different RNA-seq approaches (Hasel and collaborators (Hasel *et al.*, 2017) performed RNA-Seq of whole cell samples, while the Tabula Muris Consortium (Consortium, 2020) used single-cell RNA-Seq), we used the fold-change reported by the authors as significant differential expression and separated each group into up or downregulated after glutamatergic neurotransmission or brain aging, in each cell type.

### Mouse ortholog search for hub genes

Hub associated genes; denominated *hub genes* were originally linked to a human entrez gene ID (see above). We used the g:Profiler tool (Kolberg et al., 2020; Raudvere et al., 2019) at https://biit.cs.ut.ee/gprofiler/gost, and used the g:Orth Orthology search tool to transform human entrez gene IDs into mouse orthologs. This tool delivers the official gene symbol and Ensembl mouse IDs. The resulting mouse ortholog set was cross referenced with the hub genes set, and the intersection resulted in the four *differential hub gene* sets: 1) Differential hub genes after *glutamatergic neurotransmission* in the: a) Neuron, b) Astrocyte; and 2) Differential hub genes regulated during *brain aging* (aged/young) in the: a) Neuron, b) Astrocyte.

### KEGG Pathway enrichment analysis

The ClueGO (Bindea et al., 2009) plugin in Cytoscape (Shannon et al., 2003) was used. Mus musculus [10090] was selected, and for each subset mentioned above (1a, 1b, 2a, 2b), a separate analysis was performed, using two clusters: one for upregulated genes and the second for downregulated genes. The KEGG database from 13 May 2021 was used, the minimum number of genes per cluster was set as 2, and all other parameters were left as default. Resulting enriched KEGG pathways were manually curated to exclude terms that were unrelated to the nervous systems (see Supplemental Figures S3 to S6 for uncurated files).

## SUPPLEMENTAL INFORMATION

SUPPLEMENTAL DOCUMENT 1. Supplemental Figures S1-S6.

**Figure S1. Glucose yield is consistent with aerobic metabolism.** Phenotypic phase planes are shown as two-dimensional color maps. The Flux Balance Analysis (FBA) solution is represented by the red-filled circle. The white piecewise line depicts the specific contour level of the solution. **a.** Oxygen molecules spent per molecule of glucose. **b.** ATP molecules produced per molecule of glucose.

Figure S2. Flux coupling between sodium removal and oxidative phosphorylation in neurons.

Figure S3. Uncurated KEGG enrichment diagram for differential hub genes in the neuron during neurotransmission.

Figure S4. Uncurated KEGG enrichment diagram for differential hub genes in the neuron during brain aging.

Figure S5. Uncurated KEGG enrichment diagram for differential hub genes in the astrocyte during neurotransmission.

Figure S6. Uncurated KEGG enrichment diagram for differential hub genes in the astrocyte during brain aging.

SUPPLEMENTAL DOCUMENT 2. Supplemental Table S1.

**Table S1. Optimal fluxes relevant to the neuron-astrocyte metabolic network during neurotransmission.** The lactate shuttle is active in both directions; L-LACt2r_Int is the efflux from the astrocyte, and L-LACt2r_Neuron corresponds to the influx to neurons. Also, the glutamate-glutamine cycle was active for neuronal glutamate export (GLUVESSEC_Neuron) and glutamine efflux from astrocytes (GLNtN1_INt).

SUPPLEMENTAL DOCUMENT 3. Supplemental Theoretical Framework.

