## Supplemental document 1 for "Metabolic switch in the aging astrocyte supported via integrative approach comprising network and transcriptome analyses"

### SUPPLEMENTAL FIGURES

**Figure S1. Glucose yield is consistent with aerobic metabolism.** Phenotypic phase planes are shown as two-dimensional color maps. The Flux Balance Analysis (FBA) solution is represented by the red-filled circle. The white piecewise line depicts the specific contour level of the solution. **a.** Oxygen molecules spent per molecule of glucose. **b.** ATP molecules produced per molecule of glucose.

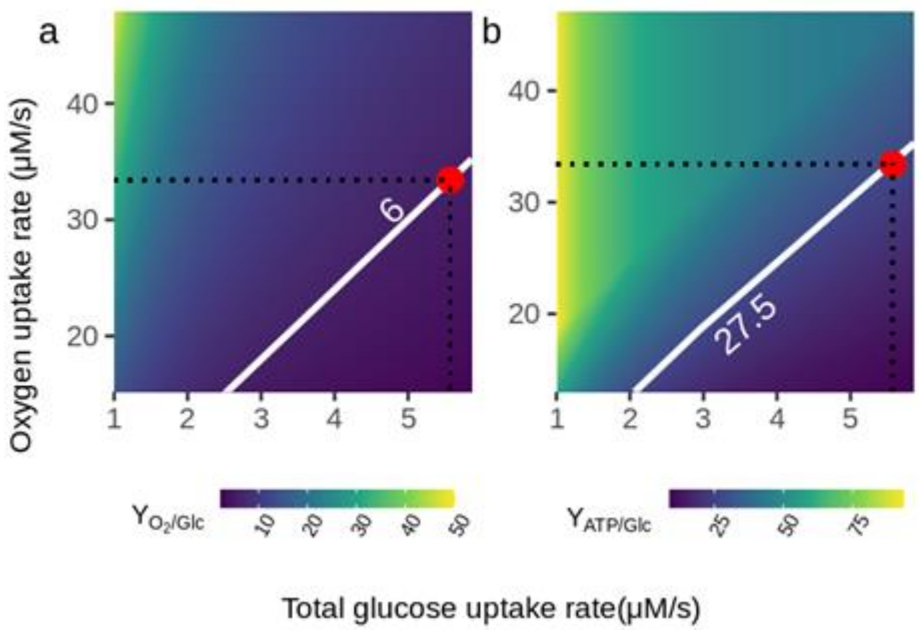

**Figure S2. Flux coupling between sodium removal and oxidative phosphorylation in neurons.**

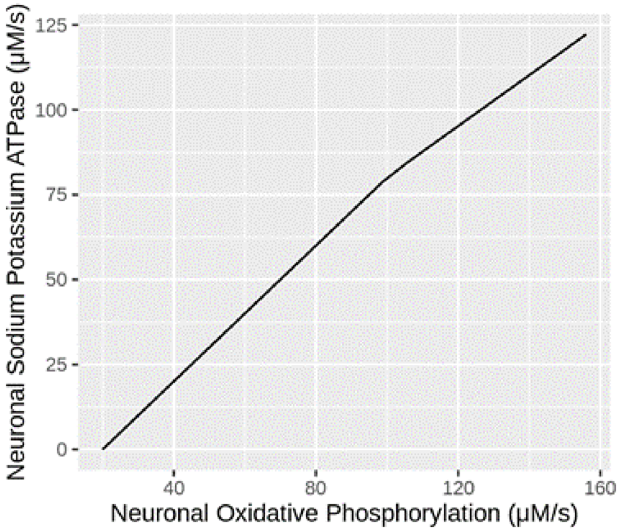

**Figure S3. Uncurated KEGG enrichment diagram for differential hub genes in the neuron during neurotransmission.**

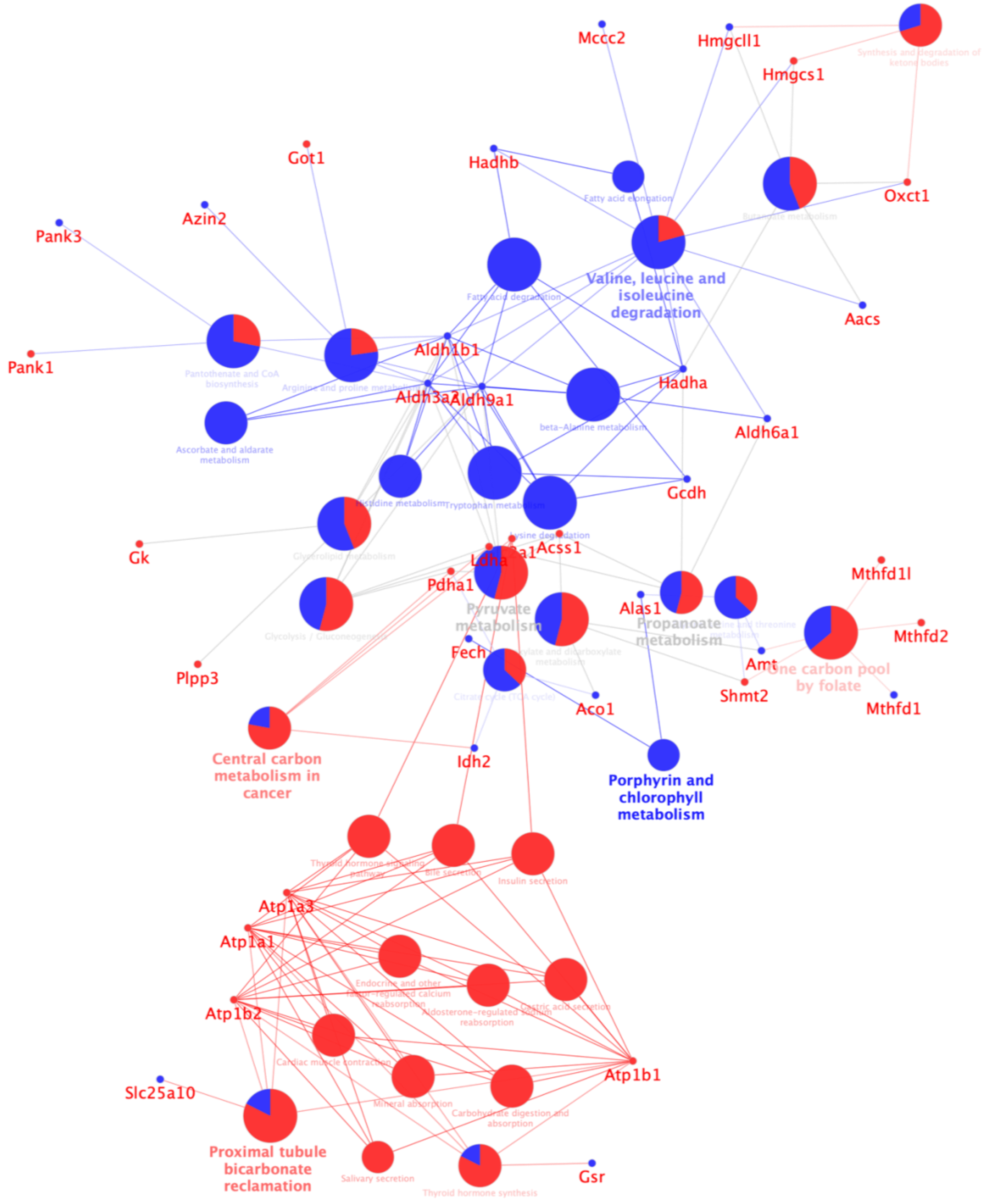

**Figure S4. Uncurated KEGG enrichment diagram for differential hub genes in the neuron during brain aging.**

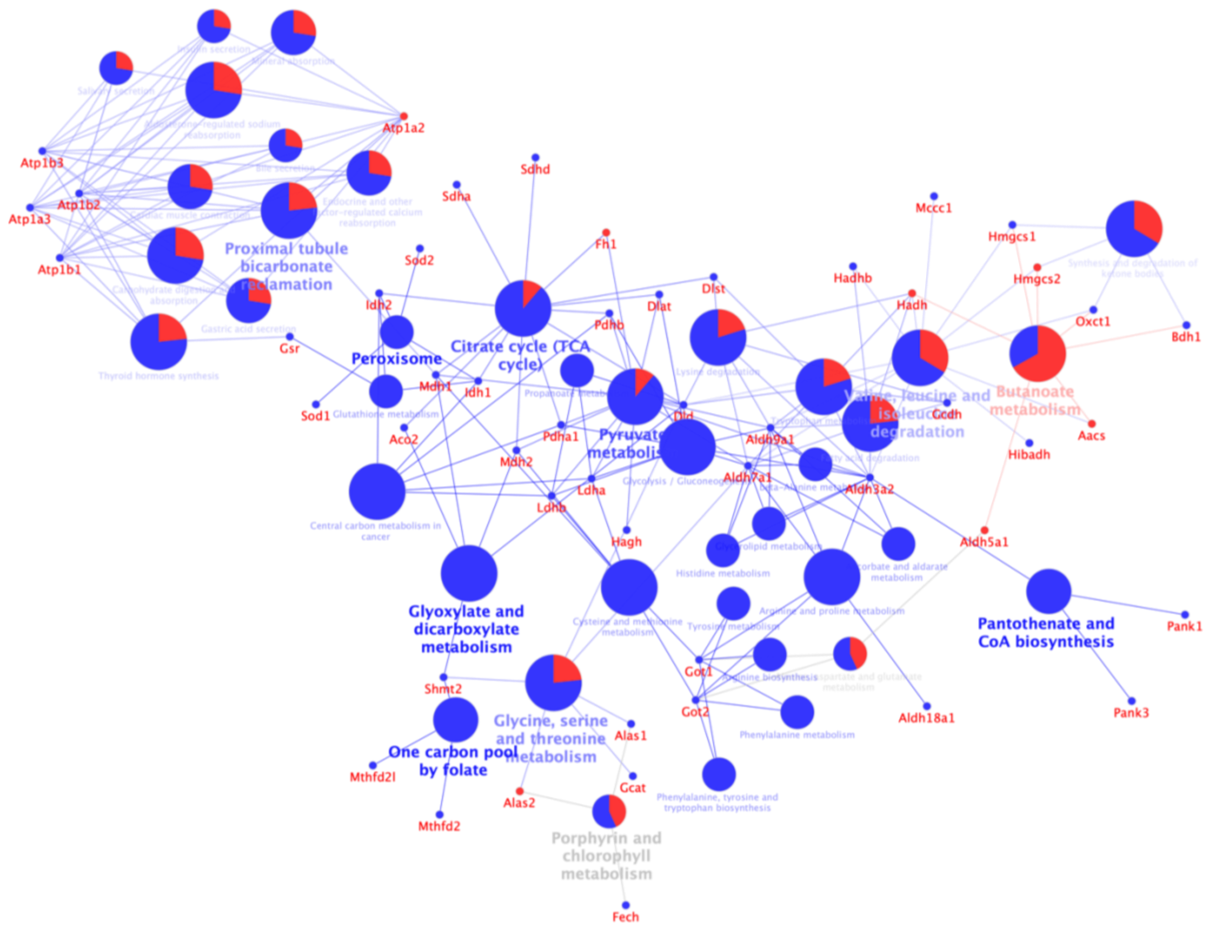

**Figure S5. Uncurated KEGG enrichment diagram for differential hub genes in the astrocyte during neurotransmission.**

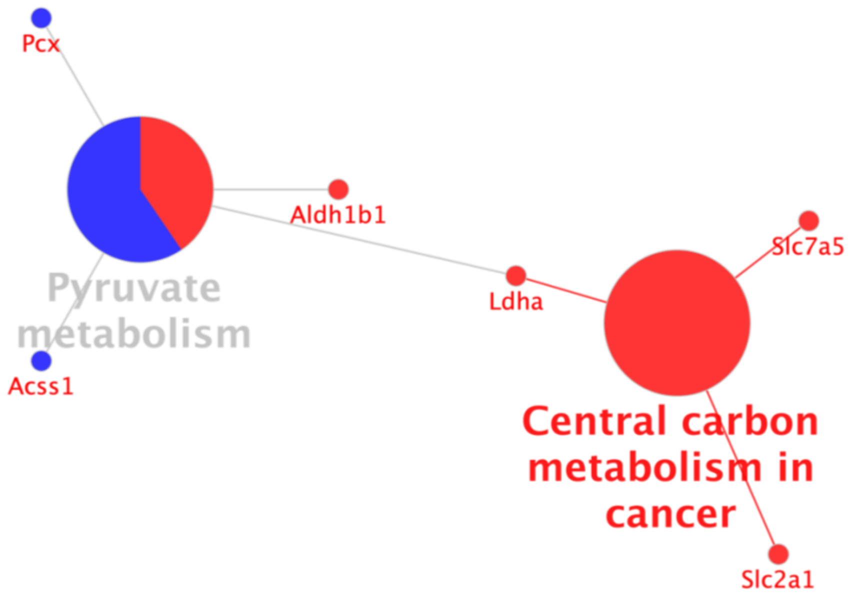

**Figure S6. Uncurated KEGG enrichment diagram for differential hub genes in the astrocyte during brain aging.**

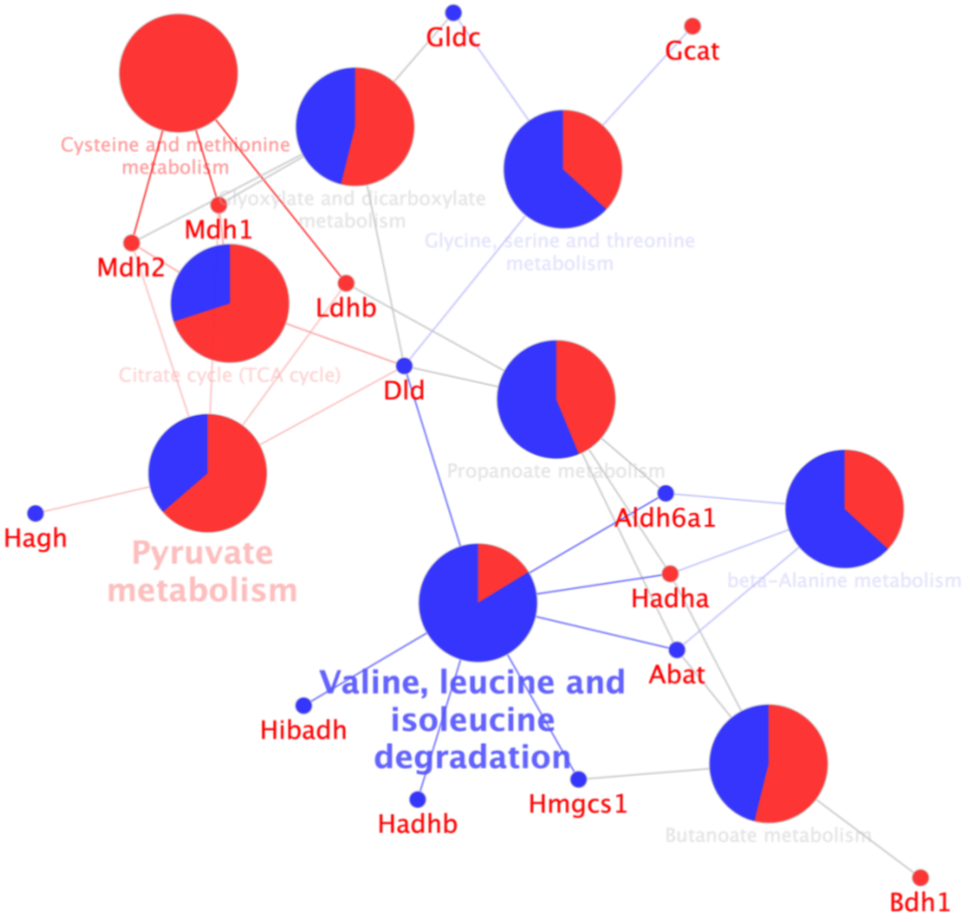
