## Supplemental document 2 for "Metabolic switch in the aging astrocyte supported via integrative approach comprising network and transcriptome analyses"

**Table S1.** Optimal fluxes relevant to the neuron-astrocyte metabolic network during neurotransmission. The lactate shuttle is active in both directions; L-LACt2r_Int is the efflux from the astrocyte, and L-LACt2r_Neuron corresponds to the influx to neurons. Also, the glutamate-glutamine cycle was active for neuronal glutamate export (GLUVESSEC_Neuron) and glutamine efflux from astrocytes (GLNtN1_INt).

| | Name | Reaction | Flux | Sensitivity |
|---|---|---|---|---|
| **EX_o2(e)** | Oxygen upkate | o2[e] <-- | -33.425896 | 0.000000e+00 |
| **NaEX_Neuron** | Neuronal sodium accumulation rate under stimulation | na1[e] <=> na1[cN] | 350.000024 | 0.000000e+00 |
| **NaKt_Neuron** | Neuronal sodium-potassium ATPase pump (sodium removal)) | atp[cN] + h2o[cN] + 2.0 k[I] + 3.0 na1[cN] --> adp[cN] + h[cN] + 2.0 k[cN] + 3.0 na1[I] + pi[cN] | 118.045878 | 0.000000e+00 |
| **L-LACt2r_Int** | L-lactate reversible transport via proton symport Interstitial And Synapse | h[I] + lac-L[I] <-- h[cA] + lac-L[cA] | -6.912674 | -0.000000e+00 |
| **L-LACt2r_Neuron** | L-lactate reversible transport via proton symport Neuron | h[I] + lac-L[I] --> h[cN] + lac-L[cN] | 6.912674 | 0.000000e+00 |
| **GLUVESSEC_Neuron** | L-glutamate secretion via secretory vesicle (ATP driven) Neuron | atp[cN] + glu-L[cN] + h2o[cN] --> adp[cN] + glu-L[I] + h[cN] + pi[cN] | 4.137608 | 0.000000e+00 |
| **GLNtN1_Int** | Glutamine transporter Interstitial And Synapse | gln-L[I] + h[cA] + na1[I] <=> gln-L[cA] + h[I] + na1[cA] | -4.137608 | -2.775558e-16 |
| **ATPS4m_Neuron** | ATP synthase (four protons for one ATP) Neuron | adp[mN] + 4.0 h[cN] + pi[mN] --> atp[mN] + h2o[mN] + 3.0 h[mN] | 155.943088 | 0.000000e+00 |
| **ATPS4m** | ATP synthase (four protons for one ATP) Astrocyte | adp[mA] + 4.0 h[cA] + pi[mA] <-- atp[mA] + h2o[mA] + 3.0 h[mA] | -0.016660 | 8.630000e+00 |
| **GLCt1r** | glucose transport (uniport) Astrocyte | glc-D[e] --> glc-D[cA] | 3.452172 | 6.652000e+01 |
| **GLCt1r_Neuron** | Glucose transporter_Neuron | glc-D[e] --> glc-D[cN] | 2.120199 | 6.000000e+01 |
| **ATPtm_Neuron** | ADP/ATP transporter, mitochondrial Neuron | adp[cN] + atp[mN] --> adp[mN] + atp[cN] | 155.943088 | 0.000000e+00 |
| **PYK_Neuron** | pyruvate kinase Neuron | adp[cN] + h[cN] + pep[cN] --> atp[cN] + pyr[cN] | 4.240398 | 0.000000e+00 |
| **PYK** | pyruvate kinase Astrocyte | adp[cA] + h[cA] + pep[cA] --> atp[cA] + pyr[cA] | 6.896014 | 0.000000e+00 |
