## Supplemental document 3 for "Metabolic switch in the aging astrocyte supported via integrative approach comprising network and transcriptome analyses"

**Supplementary Theoretical Framework**

1. Constraint-based stoichiometric modeling

We briefly present the fundamentals of constraint-based stoichiometric modeling to introduce readers from other fields. The rationale behind the constraint-based stoichiometric models of cellular metabolism (Bordbar et al., 2014) stems from fundamental physicochemical principles (Gottstein et al., 2016) that are exposed in the following sections.

1.1 Fundamental derivation of constraints-based stoichiometric models

Based on Gottstein et al., (2016), in the first place we define the concentration of metabolite $i$ as

$$c_i = n_i/V \tag{S1.1}$$

being $n_i$ the number molecules of metabolite $i$, and $V$ the volume of the cell. Taking into account that metabolite concentration undergoes dynamic changes, $n_i$ and $V$ are time-dependent. By applying the quotient rule to derive equation S1.1, the evolution over time of $c_i$ is

$$\begin{aligned} \frac{d}{dt}c_i &= \frac{d}{dt}\left(\frac{n_i(t)}{V(t)}\right) \\ &= \frac{V \times n_i'(t) - n_i \times V'(t)}{V^2} \\ &= \frac{V \times n_i'(t)}{V^2} - \frac{n_i \times V'(t)}{V^2} \\ &= V^{-1}\frac{\partial n_i}{\partial t} - n_i V^{-2}\frac{\partial V}{\partial t} \end{aligned} \tag{S1.2}$$

Assuming that neither astrocytes nor neurons change their volume during neurotransmission, we have that $\frac{\partial V}{\partial t} = 0$, then S1.2 can be rewritten as

$$\frac{d}{dt}c_i = V^{-1}\frac{\partial n_i}{\partial t} \tag{S1.3}$$

Here, $V^{-1}\frac{\partial n_i}{\partial t}$ corresponds to the net flux of metabolite $i$. Each metabolite can be produced, exchanged, or consumed by different reactions. Thus, metabolite net balance is equivalent to the flux balance of all reactions in which the metabolite participates in. Transporters and channels are considered as reactions since they also generate flux. Formally, the mass balance for metabolite $i$ consists of the sum of all fluxes ($n$ in total) multiplied by their respective stoichiometric coefficients

$$\sum_{j=1}^{n} s_{ij} v_j \tag{S1.4}$$

where $s_{ij}$ represents the stoichiometric coefficients for metabolite $i$ regarding reaction $j$. These coefficients are zero when metabolite $i$ does not participate in reaction $j$, negative for reactions that consume metabolite $i$, and positive for reactions that produce metabolite $i$. In can be noted that equations S1.3 and S1.4 are equivalent, then

$$\frac{d}{dt} c_i = V^{-1}\frac{\partial n_i}{\partial t} = \sum_{j=1}^{n} s_{ij} v_j \tag{S1.5}$$

We can generalize S1.5 for $m$ metabolites and $n$ reactions using vector notation as follows

$$\frac{d}{dt} c = Sv \tag{S1.6}$$

here, $\frac{dc}{dt}$ is an $m\,x\,1$ vector that contains the terms $\frac{dc_i}{dt}$ $(i = 1,\ldots, m)$, while $Sv$ is the vector notation for $\sum_j s_{ij} v_j$. The term $S$ is an $m\,x\,n$ matrix that contains all the stoichiometric coefficients, and $v$ is a $n\,x\,1$ vector that represents all the fluxes. In this way, equation S1.6 accounts for all of the relationships between metabolites and reactions in a given network being the cornerstone of constraint-based modeling.

#### 1.2 The dynamic steady-state assumption in the neuron-astrocyte metabolic network

The metabolic state where no accumulation or depletion of intracellular metabolites occurs is a steady state. Although metabolic networks are not strictly steady-state systems, many homeostatic states over short periods of time are close to being stationary (Fang et al., 2020; Palsson, 2015), including neurotransmission. In this sense, neuronal metabolism can adjust its fluxes to maintain ATP and ADP levels invariable (Baeza-Lehnert et al., 2019). The steady-state implies that intracellular fluxes are balanced, thus yielding zero variation in intracellular metabolite concentration. Under this condition eq. S1.6 can be rewritten as

$$Sv = 0 \qquad \text{(S1.7)}$$

Equation S1.7 can be regarded as an abstraction of a dynamic quasi-stationary metabolic state. This steady-state is dynamic because boundary metabolites can have a non-zero balance, thus allowing input and output fluxes (Fell, 2021; Yasemi and Jolicoeur, 2021). For instance, in the case of neurotransmission, extracellular glucose and oxygen are constantly depleted and replenished; thus, both molecules yield a negative (non-steady) balance. To maintain the right-hand side of equation S1.7 as a zero column vector, extracellular glucose and oxygen mass balances are composed of two balanced fluxes, one associated with the uptake (negative flux) and the other simulating the "incoming" of new extracellular molecules (positive flux). The same principle is applied to secreted metabolic products that have positive (non-steady) balance, such as the lactate produced by the astrocyte. In this case, the sense of the fluxes is in the opposite direction of the substrates, and the flux that simulates the "incoming" of new extracellular molecules is replaced by a flux that accounts for the "wash out" of extracellular molecules. Biologically, a dynamic steady-state occurs when constant "incoming" and "wash out" of extracellular molecules can be assumed, avoiding substrate exhaustion and product accumulation in the cell. Metabolites associated with constant "incoming" and "wash out" are boundary metabolites. In the neuron-astrocyte metabolic network, glucose, oxygen, lactate, glutamate, glutamine, and sodium ions are boundary metabolites and define the response to neurotransmission workload. Hence, we assumed a constant replenishment of oxygen and glucose, thus ensuring that boundary uptake fluxes are always established. Also, we considered that lactate, glutamate, and glutamine are rapidly exchanged between neurons and astrocytes, emulating an extracellular "wash out." Finally, we regarded extracellular sodium ions as fast-diffusing ions that rapidly diffuse.

#### 1.3 The null space of $S$ and flux constraints

Given the degrees of freedom the model has (more reactions than metabolites), fluxes can vary with no fixed ratios, and the system still would be at steady-state. This means that vector $v$ in equation

S1.7 represents a set of flux vectors that satisfy the steady-state condition. Mathematically, this set of vectors is known as the null space of $S$, and contains all the stationary flux distributions attainable by the model (Palsson, 2015). In this sense, equation S1.7 is regarded as the steady-state constraint. Flux direction (that accounts for reaction thermodynamics) and flux bounds (lower and upper limits) also fall into the category of constraints and correspond to inequalities expressed as

$$L_b \leq v \leq U_b \tag{S1.8}$$

where $v$ is the flux vector while $L_B$ and $U_B$ are the lower, and upper bounds, respectively. These conditions plus equation S1.7 are constraints that define a space of attainable metabolic states, also known as the feasible space. All constraint-based models are formulated based on these principles.

#### 1.4 Computing biologically meaningful metabolic states

Constraint-based models have many possible metabolic states, which are contained in the null space of $S$. We can assume that any biologically meaningful state is optimized for whatever the cell is aiming to achieve. Hence, physiologically states may be predictable by employing optimization-based approaches, such as linear programming (Heirendt et al., 2019). In order to determine this biologically meaningful metabolic state, optimality conditions should be imposed. In this sense, we define an objective function which encodes what is aimed to be optimized. Objectives are context-specific and depend on the particular phenotype or cellular compartment that one tries to model (Chen et al., 2019; Feist and Palsson, 2010; Sánchez et al., 2012; Smith and Robinson, 2011). The steady-state and flux constraints serve as constraints for the optimization. Once solved, the solution is associated with an optimal metabolic state, or optimal flux distribution. This optimization approach is known as Flux Balance Analysis (Orth et al., 2010) and is formulated as

$$\begin{aligned} &\text{Maximize } z = c^T v \\ &\text{Subject to} \begin{cases} Sv = 0 \\ L_b \leq v \leq U_b \end{cases} \end{aligned} \tag{S1.9}$$

here, the objective function is $c^T v$, which correspond to a linear combination of the fluxes that are involved in the objective.

### 2. Analysis based on network topology

This section explains the theoretical basis for the topological analysis of the stoichiometric matrix and shows how such an approach complements what the FBA informs. Readers already introduced to the mathematics of constraint-based modeling may skip all previous sections. However, for readers outside the field, we highly recommend them revising this supplementary material from the beginning since this section builds upon previous ones.

#### 2.1. The stoichiometric matrix as a bipartite network

An intuitive network representation of the metabolism is a bipartite network. A bipartite network has two types of nodes with edges connecting only nodes from the different types. For metabolic networks, the two types of nodes represent metabolites and reactions, with edges joining each metabolite to the reactions it participates in (either as a product or substrate). The algebraic representation of a bipartite metabolic network is an incidence or bi-adjacency matrix. Considering *n* as the number of reactions and *m* as the number of metabolites, the incidence matrix $B$ is a *m* x *n* matrix that has elements $B_{ij}$ such that

$$B_{ij} = \begin{cases} 1 & \text{if reaction } j \text{ uses or produces metabolite } i \\ 0 & \text{otherwise} \end{cases} \qquad \text{(S2.1)}$$

In our case, the stoichiometric matrix $S$ can be transformed into the matrix $B$ by binarizing it. In this sense, let us denote any given element of $S$ as $s_{ij}$, then denote the binary version of $S$ as $\widehat{S}$. The elements of $\widehat{S}$ are computed as

$$\widehat{s_{ij}} = \begin{cases} 1 & \text{if } s_{ij} = 0 \\ 0 & \text{if } s_{ij} \neq 0 \end{cases} \qquad \text{(S2.2)}$$

Here, if $s_{ij}$ is one, the metabolite $i$ participates in reaction $j$. It can be noted that $\widehat{S}$ is equal to $B$. As we will see in the following sections, the matrix $\widehat{S}$ is key to perfomed the topological-based analysis.

#### 2.2 Reaction projection of the stoichiometric matrix

As we are interested in finding critical genes, hence enzymes and transporters, we constructed the adjacency matrix for the relationships between reactions. This reaction adjacency matrix was computed from column projection of $\hat{S}$ as follows

$$A_v = \widehat{S^T}\widehat{S} \tag{S2.3}$$

Here, the off-diagonal elements of the matrix $A_v$ are the inner product of the columns of $\hat{S}$, i.e. $(a_v)_{ji} = \hat{s}_j^T \hat{s}_i$. Each of the off-diagonal elements indicates the number of metabolites that reactions *i* and *j* have in common. The diagonal elements of $A_v$ correspond to the number of metabolites that participate in each reaction. Because we are interested in the fundamental structure of the network, $A_v$ was binarized and its diagonal was set to zero. Finally, we denoted this last binary adjacency matrix as $A$, and was used for all of the centrality calculations. Figure S3 illustrates the main steps for deriving the reaction (binary) adjacency matrix from the stoichiometric matrix.

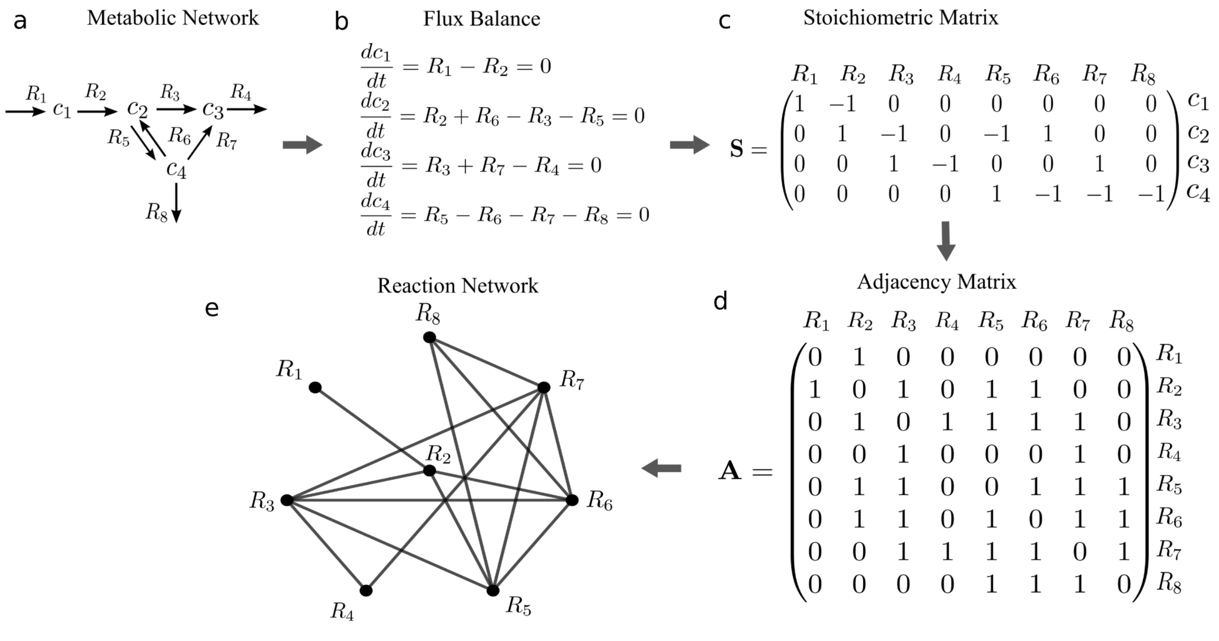


d°

**Figure S3.** Projection of the stoichiometric matrix into a binary adjacency matrix of reactions.

2.3 Node centrality as an index of node importance and availability

Centrality metrics are a group of topology-based measures that quantify the relevance of a node in a network. These metrics inform the level of integration or availability a node has in the network by encoding different aspects of its topological context (Borgatti and Everett, 2021). High availability means that a node can influence or be influenced by others, thus informing about its integration. In our work, we combined two different kinds of centrality metrics to quantify the availability of the sensitivity nodes previously computed via FBA. One kind of centrality was a proxy of the probability of having an interaction with any given node, and the other kind represented the inverse of the cost of establishing such an interaction. Thus, our index for nodal availability was formulated so that high availability may be reached by having a high probability of receiving signals, having easy access, or both. Such a probability was computed using a degree-like centrality metric, while the cost was estimated by using closeness-like centrality metrics. In simple terms, degree-like centralities count the trajectories in which a node participates, while closeness-like centralities measure the length of such trajectories. Thus, degree-like metrics encode the number of possible interactions (probability), and closeness-like metrics encode the node's reachability (cost). In graph-theoretical terms, these two metrics are called *radial centralities* because they consider the node in question as the endpoint of interactions (Borgatti and Everett, 2006). Thus, the sum of the two metrics yields the radial involvement of the node. Henceforward, we will adopt the state-of-the-art denominations for the centrality metrics; namely, degree-like centrality will be referred to as connectedness-based centrality and closeness-like as closeness-based centrality (Borgatti and Everett, 2021). We used one connectedness-based and two closeness-based centralities which at the end were aggregated into a single quantity of node centrality. In the following section, we provide the mathematical formulations of these metrics.

### 3.0 Centrality metrics description

#### 3.1 Connectedness-based centrality

Connectedness-based centralities count the unrestricted trajectories in which a node participates as the ending point. Notably, these metrics represent the certainty or probability of receiving a signal.

These centralities are computed by employing the eigendecomposition of the adjacency matrix (Newman, 2018).

***Eigenvector centrality (EC).*** This centrality metric accounts for the quantity and quality of a node's connections by accounting for its degree and the degree of its neighbors (Bonacich, 1987; Fornito et al., 2016). The eigenvector centrality of node *i* ($EC_i$) is proportional to the sum of the centralities of its neighbors. The eigenvectors ($x$) and eigenvalues ($\lambda$) of the adjacency matrix ($A$) are employed for the calculation, where the eigendecomposition $Ax = \lambda x$ is rearranged. Having $A$ as our adjacency matrix, the eigenvector centrality of reaction *i* is defined as

$$EC_i = x_i = \frac{1}{\lambda_1} \sum_{j=1}^{n} A_{ij} x_j \quad \text{(S3.1)}$$

Here, $\lambda_1$ is the leading eigenvector, i.e., the eigenvector corresponding to the largest (positive) eigenvalue (Fornito et al., 2016; Newman, 2018).

3.2 Closeness-based centralities

Two nodes are topologically close if a path with few edges connects them. In this sense, a node has high closeness centrality if, on average, it is topologically close to many other nodes. A node with a short average path length can interact with many network elements via only a few links, meaning that it is topologically central (Fornito et al., 2016). Closeness-based centralities are different from the eigenvector-derived (connectedness-based) ones as they measure the trajectories' length instead of counting them.

***Closeness centrality (CC).*** This centrality measures the mean distance from a node to other nodes. Let $d_{ij}$ be the shortest distance from node $i$ to $j$, then the mean distance from $i$ to every node is

$$l_i = \frac{1}{n-1} \sum_{j=1}^{n} d_{ij} \quad \text{(S3.2)}$$

then, the CC for node $i$ is (Beauchamp, 1965):

$$CC_i = \frac{1}{l_i} = \frac{n-1}{\sum_{j=1}^{n} d_{ij}} \quad \text{(S3.3)}$$

***Information centrality (IC).*** This metric makes use of all paths between pairs of nodes but gives them relative weighting as a function of the information they transmit (Stephenson and Zelen, 1989). This kind of centrality differs from closeness centrality as it does not only consider shortest paths. The IC assumes that all paths do not transmit the same information. If nodes i and j are connected by $k_{ij}$ paths, such paths are

$$P_{ij}(s) \text{ for } s = 1 \ldots k_{ij} \tag{S3.4}$$

for each path we have

$$I_{ij}(s) = \frac{1}{D_{ij}(s)} \tag{S3.5}$$

Where $D_{ij}(s)$ is the length of $P_{ij}(s)$ from equation S3.5, the resulting quantity $I_{ij}(s)$ is defined as the information of $P_{ij}(s)$. Then we combine the information of the paths between the  nodes i and j

$$C_{ij} = \sum_{s=1}^{k_{ij}} l_{ij} \tag{S3.6}$$

and finally, we compute the information centrality for node $i$ as the harmonic mean of the combined information of the paths from $i$ to all the other nodes

$$I_i = \frac{n}{\sum_{s=1}^{j_{ij}} C_{ij}^{-1}} \tag{S3.7}$$

(Stephenson and Zelen, 1989).

### 4.0 Reactions topology can represent metabolic states outside the steady state

We can interpret equation S1.6 as the row space of $S$. According to the properties of the four fundamental subspaces of linear transformations, the row space is orthogonal to the null space, which means that the row space contains the non-steady state mass balances of $S$ (Palsson, 2015). Thus we can express the row space of $S$ as

$$Sv_{\text{dynamic}} = b_{\text{non-steady}} \tag{S4.1}$$

where $v_{dynamic}$ is a dynamic flux distribution, and $b_{non-steady}$ a metabolite mass balance outside steady-state. We can express $v_{dynamic}$ as

$$\begin{aligned} S^T S v_{\text{dynamic}} &= S^T b_{\text{non-steady}} \\ v_{\text{dynamic}} &= (S^T S)^{-1} S^T b_{\text{non-steady}} \\ v_{\text{dynamic}} &= A^{-1} S^T b_{\text{non-steady}} \end{aligned} \tag{S4.2}$$

here, $A$ is invertible as it is a squared matrix, and it is equivalent to the product shown in equation S2.3, then it encodes the reactions topology. This shows that any given non-steady metabolic state can be represented as a function of the reactions topology. The fact that reaction topology represents non-steady states allows the use of centrality metrics to determine reaction importance in states that are not considered by the FBA. Thus making centrality analysis and FBA complementary approaches.

### 5.0 Proof of concept of the entire network analysis

The whole network analysis tackled two aspects of network functionality, one regarding flux-related behavior and the other related to topology, where the former yields two kinds of results, i.e., fluxes and sensitivities. Flux informs about execution and the latter about control. Reactions with high sensitivity values were categorized into the so-called sensitivity set. This group acts as an "interface" through which the whole network can perturb the optimal response to metabolic workload. Even though both quantities (flux and sensitivity) successfully described the behavior of the neuron-astrocyte network during neurotransmission, these are limited to model only stationary fast responses. Hence, we formulated another quantity that measures each node's controllability over the centrality of the nodes belonging to the sensitivity set, we named such a quantity as Absolute Centrality Contribution (ACC). In other words, we measured how much any given node controls the level of integration or availability of the sensitivity set or “interface”. The nodes with substantial control over the availability of the interface can be regarded as gatekeepers. In sum, our approach had three outcomes: fluxes, sensitivities, and absolute centralities. In Figure S4, we provide a proof-of-concept for our approach, where a small-scale pseudo-metabolic network was subjected to the complete network-analysis workflow. Here, fluxes were computed using the objective pathway P5 in Figure S4a, thus obtaining the optimal metabolic response (Figure S4d). Sensitivities were also computed, and the sensitivity set comprised reactions with non-zero sensitivity, i.e., ATP_sink, A_uptake, and L_diffusion. These reactions constitute the "interface" (Figure S4e). Then we computed the ACC for each node and determined the nodes with the highest AC. For simplicity, we denoted the top 3 ACC reactions as the central metabolic reactions or gatekeepers (Figure S4f).

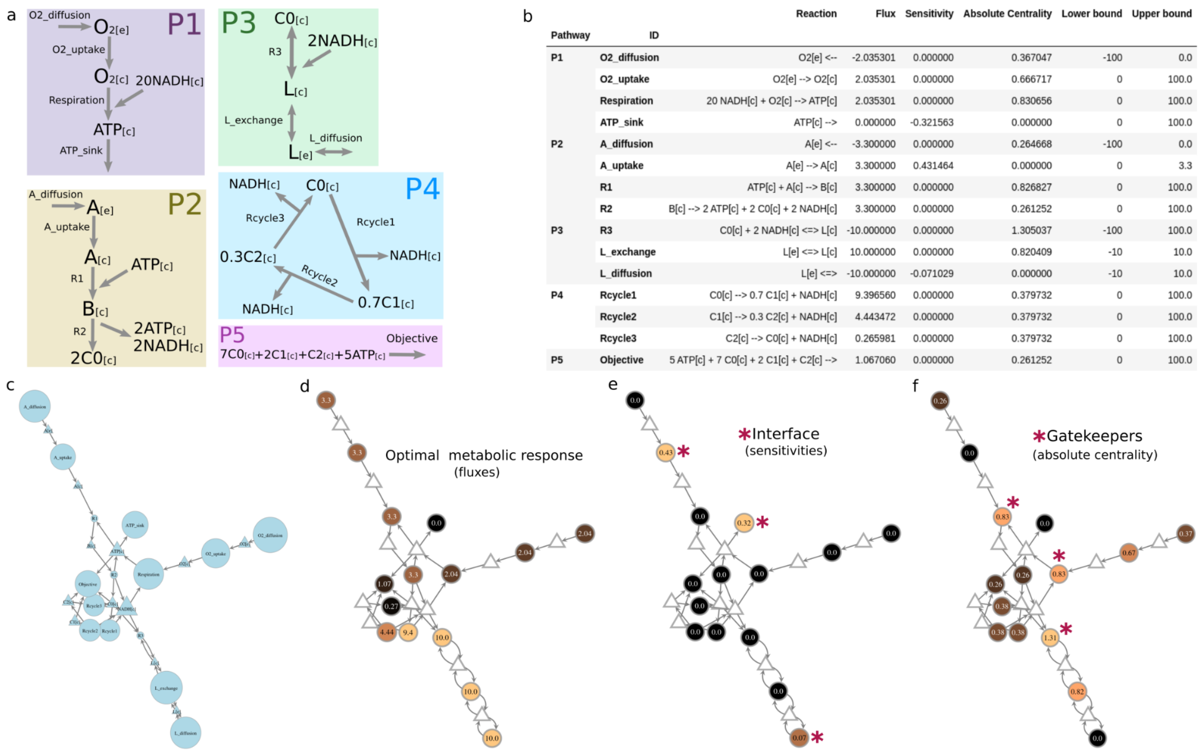


| Pathway | ID | Reaction | Flux | Sensitivity | Absolute Centrality | Lower bound | Upper bound |
|---|---|---|---|---|---|---|---|
| P1 | O2_diffusion | O2[e] <-- | -2.035301 | 0.000000 | 0.367047 | -100 | 0.0 |
| | O2_uptake | O2[e] --> O2[c] | 2.035301 | 0.000000 | 0.666717 | 0 | 100.0 |
| | Respiration | 20 NADH[c] + O2[c] --> ATP[c] | 2.035301 | 0.000000 | 0.830656 | 0 | 100.0 |
| | ATP_sink | ATP[c] --> | 0.000000 | -0.321563 | 0.000000 | 0 | 100.0 |
| P2 | A_diffusion | A[e] <-- | -3.300000 | 0.000000 | 0.264668 | -100 | 0.0 |
| | A_uptake | A[e] --> A[c] | 3.300000 | 0.431464 | 0.000000 | 0 | 3.3 |
| | R1 | ATP[c] + A[c] --> B[c] | 3.300000 | 0.000000 | 0.826827 | 0 | 100.0 |
| | R2 | B[c] --> 2 ATP[c] + 2 C0[c] + 2 NADH[c] | 3.300000 | 0.000000 | 0.261252 | 0 | 100.0 |
| P3 | R3 | C0[c] + 2 NADH[c] <=> L[c] | -10.000000 | 0.000000 | 1.305037 | -100 | 100.0 |
| | L_exchange | L[e] <=> L[c] | 10.000000 | 0.000000 | 0.820409 | -10 | 10.0 |
| | L_diffusion | L[e] <=> | -10.000000 | -0.071029 | 0.000000 | -10 | 10.0 |
| P4 | Rcycle1 | C0[c] --> 0.7 C1[c] + NADH[c] | 9.396560 | 0.000000 | 0.379732 | 0 | 100.0 |
| | Rcycle2 | C1[c] --> 0.3 C2[c] + NADH[c] | 4.443472 | 0.000000 | 0.379732 | 0 | 100.0 |
| | Rcycle3 | C2[c] --> C0[c] + NADH[c] | 0.265981 | 0.000000 | 0.379732 | 0 | 100.0 |
| P5 | Objective | 5 ATP[c] + 7 C0[c] + 2 C1[c] + C2[c] --> | 1.067060 | 0.000000 | 0.261252 | 0 | 100.0 |

**Figure S4.** Network-analysis workflow applied on a small-scale network mimicking metabolic pathways. **a:** pseudo-pathways representing different metabolic subsystems: respiration and oxidative phosphorylation (P1), glycolysis (P2), lactate exchange (P3), Krebs Cycle (P4), and an energy-demanding metabolic objective (P5). **b**: results from the network-analysis workflow. **c**: bipartite network representation of the network; here, circles correspond to reactions, and triangles are metabolites. Node size was adjusted to fit the text inside. **d**: Fluxes computed by maximizing the P5 pathway. **e**: sensitivities, non-zero sensitivities are considered as the interface. **f**: absolute centrality values, where the top 3 were classified as central metabolic reactions or gatekeepers.

Steady-State and Kinetic Approaches. Processes *9*, 322. https://doi.org/10.3390/pr9020322.
